# In the words of others: ERP evidence of speaker-specific phonological prediction

**DOI:** 10.1101/2025.04.16.648895

**Authors:** Marco Sala, Francesco Vespignani, Simone Gastaldon, Laura Casalino, Francesca Peressotti

**Author notes:** Corresponding authors: Marco Sala Francesca Peressotti Dipartimento di Psicologia dello Sviluppo e della Socializzazione Università degli Studi di Padova Via Venezia 8, 35131 Padova (PD), Italy.

## Abstract

Prediction models usually assume that highly constraining contexts allow the pre-activation of phonological information. However, the evidence for phonological prediction is mixed and controversial. In this study, we implement a paradigm that capitalizes on the phonological errors produced by non-native speakers to investigate whether speaker-specific phonological predictions are made based on speaker identity (native-vs-foreign). EEG data was recorded from 42 healthy native Italian speakers. Participants were asked to read sentence fragments after which a final word was spoken by either a native- or a foreign-accented speaker. The spoken final word could be predictable or not, depending on the sentence context. The identity of the speaker (native-vs-foreign) may or may not be cued by an image of the face of the speaker. Our main analysis indicated that cueing the speaker identity was associated with a larger N400 predictability effect, possibly reflecting an easier processing of predictable words due to phonological preactivation. As visual inspection of the waveforms revealed a more complex pattern than initially anticipated, we used temporal EFA (Exploratory Factor Analysis) to identify and disentangle the ERP components underlying the effect observed. In the native-accent condition, predictable words elicited a posterior positivity relative to unpredictable words, possibly reflecting a P3b response, which was more pronounced when the speaker identity was cued. In the foreign-accent condition, cueing the speaker identity was associated with a smaller N1 and a larger P3a response. These results suggest that phonological prediction for native- and foreign-accented speakers likely involve different cognitive processes.

## Introduction

In both oral and written language, the intended meaning of a sentence emerges as more information becomes available. The linearity of the linguistic input suggests that complex meanings are constructed gradually, as words are processed: the meaning of each word is retrieved from the continuous sensory input and integrated into an unfolding representation of the preceding context (Altmann & Mirković, 2009; Frazier & Rayner, 1982). Nevertheless, compelling evidence from different cognitive domains has shown that the human brain predicts incoming information to optimize information processing (Cheung & Bar, 2012; Costa et al., 2024; Friston, 2005).

Prominent models of language comprehension propose that highly constraining contexts enable the preactivation of meaning and form of the expected linguistic input (Altmann & Mirković, 2009; Dell & Chang, 2014; Kuperberg & Jaeger, 2016; Pickering & Gambi, 2018; Pickering & Garrod, 2007, 2013). Consistent evidence has been provided for the preactivation of both semantic (Altmann & Kamide, 1999, 2007; Chambers et al., 2002; Federmeier & Kutas, 1999a; Kamide et al., 2003; Metusalem et al., 2012; Paczynski & Kuperberg, 2012) and syntactic (Crocker, 2000; Kimball, 1975; Levy, 2008; Lewis, 2000; Staub & Clifton, 2006; Traxler, 2014; Traxler et al., 1998; van Gompel et al., 2005) information during sentence processing. However, the extent to which comprehenders can predict phonological forms in highly constraining contexts remains a matter of debate, with current models of language comprehension remaining rather underspecified in this regard. In the event-related potentials (ERP) literature, most studies investigating whether sentence contexts are used to predict the phonological form of a highly predictable word have focused on the N400 component (DeLong et al., 2005, 2019, 2021; Ito et al., 2016, 2017, 2020; Martin et al., 2013; Nieuwland et al., 2018). The N400 is a negative-going, centro-parietally distributed component of the ERP, peaking around 400 ms after word onset and strongly associated with lexical and semantic processing (Kutas & Federmeier, 2011; Kutas & Hillyard, 1980, 1984). The N400 is typically elicited by open-class words (Kutas & Van Petten, 1988) and smaller for high frequency words and for words which have been semantically primed. Within sentences, the N400 amplitude is highly correlated with an offline measure of the eliciting word’s predictability, known as *cloze probability* - the proportion of individuals which continue a sentence fragment with a given word - where higher cloze probability results in a smaller N400 amplitude (Kutas & Federmeier, 2011) Two main proposals have been made regarding the cognitive processes underlying the modulation of the N400 amplitude. The *integration view* hypothesizes that facilitatory effects of context arise only when the features of a verbal input have already been accessed through bottom-up processing, reflecting the degree of (mis)match between the context features and the current word features and/or an easier process of linking the current word with prior context information (Kutas & Federmeier, 2011; Van Petten, 1993; Van Berkum et al., 1999; Van Petten & Luka, 2012). The *prediction view* posits that facilitatory effects of context emerge due to the top-down preactivation of upcoming words, resulting in an easier retrieval of the corresponding lexical-semantic representation from long-term memory (DeLong et al., 2005; Federmeier & Kutas, 1999b; Kuperberg, 2016; Kutas & Federmeier, 2011; Lago et al., 2023). The two views are not incompatible with each other and some researchers argue that the N400 does not reflect a single process but rather the combined activity of multiple processes, indexing both the preactivation of the upcoming linguistic information and its integration in the sentence context (Baggio, 2012, 2018; Baggio & Hagoort, 2011; Kutas & Federmeier, 2011; Newman R. L. et al., 2012; Pylkkänen & Marantz, 2003). Emerging evidence suggests that these mechanisms may operate in parallel, contributing to similar but distinct subcomponents of the N400 (Nieuwland et al., 2020). One issue within the prediction view is about the format of these predictions. Without specifying whether the predictions involve concepts, parts of speech or expected grammatical structures, specific lexical entries, phonological or orthographic forms, the proposal remains rather generic and difficult to falsify. For this reason, empirically demonstrating the anticipated activation of specific representations has significant theoretical implications.

One of the most compelling direct empirical evidence of phonological prediction was reported by DeLong et al. (2005). Their experiment leveraged the English phonological rule in which the indefinite article appears as “*a”* before consonant-initial words and “*an”* before vowel-initial words. Participants read sentences with varying levels of contextual constraint that led to expectations of either a consonant- or vowel-initial word. The authors examined the modulation of the N400 amplitude elicited by the article, which could either match or mismatch the phonological form of the predicted noun. Results showed that N400 amplitude decreases as a function of increasing *cloze probability*, both for the target noun and, critically, for the preceding article. This suggests that participants predicted the phonological form of the upcoming word, leading to increased negativity when the article mismatched the expected form. Despite the theoretical significance of the DeLong et al. (2005) findings, subsequent attempts to replicate the N400 modulation on the pre-target article have yielded mixed results (Ito et al., 2017, 2020; Martin et al., 2013; Nicenboim et al., 2020; Nieuwland et al., 2018).

Laszlo & Federmeier (2009) investigated the prediction of phonological/orthographic word forms using stimuli phonologically/orthographically related to the predictable word. They found that the N400 amplitude was larger for unexpected words and non-words compared to predictable words. Crucially, the N400 effect was reduced for form-related words and non-words relative to form-unrelated words and non-words. Similarly, Ito et al. (2016) used a paradigm in which participants read sentences ending with either a predictable word (e.g., book), a form-related word (e.g., hook), a semantically-related word (e.g., page), or an unrelated word (e.g., sofa). They replicated the reduced N400 effect for form-related words, but this effect was found only for high-cloze sentence continuations and slow presentation rates (700 ms per word). The authors interpreted their results as supporting *prediction-by-production* accounts of linguistic prediction. Prediction-by-production models propose that prediction during comprehension relies on the same representations and mechanisms as those used in language production (Federmeier, 2007; Gastaldon et al., 2024; Huettig, 2015; Pickering & Gambi, 2018; Pickering & Garrod, 2007, 2013). According to this view, comprehenders covertly imitate what they hear, generating production-based representations of upcoming speech (Pickering & Garrod, 2013). This prediction mechanism involves a hierarchical preactivation of linguistic representations, from semantic to phonological forms as in language production (Levelt, 1999). Notably, phonological word form preactivation occurs only when sufficient time and cognitive resources are made available by the specific task and paradigm. DeLong et al. (2019, 2021) employed the same experimental design as Ito et al. (2016), but with different materials, and reported a reduced N400 effect for form-related words at both 500 and 700 ms per word presentation rates. They interpreted their findings as suggesting that phonological prediction does not necessarily require production-like processing, which is constrained by timing and resources. Instead, preactivation of upcoming words could arise from passive *spreading of activation* between linguistic representations (Anderson, 1983; Collins & Loftus, 1975; Huettig et al., 2022; Hutchison, 2003; McRae et al., 1997). Within this framework, linguistic representations can partially activate networks of related items (semantically, associatively, or phonologically; Pickering & Gambi, 2018). This mechanism is thought to be less constrained by time and/or processing resources and largely independent of the specific linguistic context, although some accounts suggest it may still be sensitive to contextual details (Huettig et al., 2022). Importantly, the proposal that spreading of activation can lead to phonological form preactivation is not incompatible with prediction-by-production theories. Pickering & Gambi (2018) proposed that linguistic preactivation could rely on both *prediction-by-production*, which is cognitively demanding and optional, and *prediction-by-association*, which is automatic but less precise. The findings of DeLong et al. (2019, 2021) suggest that phonological form preactivation can occur through mechanisms outside the language production network, or that phonological form preactivation within a prediction-by-production model may occur more rapidly than previously thought. Further studies have reported ERP (and ERF, event-related fields, for MEG) predictability effects preceding the N400, which have been interpreted as evidence of the preactivation of sublexical representations. However, their replicability remains weak and inconsistent (for a review, see Nieuwland, 2019).

Finally, phonological prediction has also been explored using behavioral techniques, such as the *visual word paradigm* in eye-tracking studies. These studies examined whether a phonological competitor of a predictable target word attracts more anticipatory looks than phonologically or semantically unrelated distractors. Results from this paradigm have been inconsistent across studies (Ito, 2019; Ito et al., 2018; Ito & Husband, 2017; Ito & Sakai, 2021; Kukona, 2020; X. Li et al., 2022; X. Li & Qu, 2024; Zhao et al., 2024). Nonetheless, a recent metanalysis (Ito et al. 2024) reported a small but reliable phonological competitor effect. In a previous behavioral study (Sala et al., 2024), we investigated whether speaker identity (native vs. foreign-accented) influences phonological predictions. Participants were first familiarized with a native and a foreign-accented speaker. Then they silently read written sentence frames that were either highly or weakly constraining towards an upcoming spoken target word, which was pronounced either by the native- or the foreign-accented speaker. The foreign-accented speaker made consistent phonological errors on the first phoneme of the target word. Crucially, during trial presentation, speaker identity was either cued or not with an image of the speaker’s face. Participants performed a lexical decision task on the spoken target word and in the foreign-accented condition, they were explicitly instructed to accept mispronounced words as correct. Our results showed that cueing the speaker identity led to faster RTs (response times) for predictable words but not for unpredictable ones, suggesting that participants used the face cue to implement phonological predictions.

The present study aims to investigate this face-cueing effect using ERPs. We adapted the experimental procedure of the previous behavioral study to the ERP context. We hypothesized that the face cue will facilitate phonological predictions, aiding phonological encoding of the target word and consequently the retrieval of the corresponding lexical-semantic representation. Therefore, cueing speaker identity should be associated with a greater N400 predictability reduction compared to when the speaker identity is not cued, reflecting easier processing of predictable words due to phonological preactivation. Additionally, we aim to explore whether the use of speaker’s accent information in prediction differs between the native- and the foreign-accented conditions. In the native-accented condition, participants should anticipate a standard phonology whereas in the foreign-accented condition they should be able to anticipate unusual specific phonological deviations.

## Methods

### Participants

A total number of 48 healthy participants (39 women) took part in the study, aged between 18 and 30 years old, with a mean age of 23.27 ± 3.05 years, and with a mean education level of 16.33 ±1.94 years. Participants were recruited from healthy volunteers and students at the University of Padova and received 15 euros for their participation. Participants were right-handed native Italian speakers with no history of neurological, language-related, or psychiatric disorders. The research adhered to the principles outlined in the Declaration of Helsinki. Participants provided their informed consent before participating in the experiment. The research protocol was approved by the Ethics Committee for Psychological Research of the University of Padova (protocol number: 5181).

### Materials

The materials will be made available on OSF upon acceptance of the manuscript. The target stimuli consisted of 168 spoken words (mean length = 5.86 ± 1.86 phonemes) beginning with the phonemes /r/, /p/ and /k/. These three phonemes were not present in any other position within words. Each target word was preceded by a written sentential context that could be either High Constraining (HC, example in 1a) or Low Constraining (LC, example in 1b) towards the target word.

1a) La carota è il cibo preferito dei conigli

The carrot is the food preferred by-the rabbits

‘The carrot is the rabbits’ favorite food’

1b) Al parco abbiamo visto una famiglia di conigli

At-the park have seen a family of rabbits

‘At the park we saw a family of rabbits’

To determine the constraint level, an online sentence completion questionnaire was administered to 22 participants, who did not take part in the experiment. They were instructed to complete each sentence frame with the first word that came to their mind. The sentence constraint was operationalized as the proportion of total responses involving the most frequent continuation (*High Constraint*: mean _sentence constraint_ = 0.94 ± 0.07; *Low Constraint*: mean _sentence constraint_ = 0.17 ± 0.09). The target word in HC sentence frames was always the most frequent continuation (*High Constraint*: mean _cloze probability_ = 0.94 ± 0.07), while in LC sentence frames it was a semantically plausible continuation (*Low Constraint*: mean _cloze probability_ = 0.03 ± 0.09). The sentence frames varied in length (mean = 9.37 ± 2.11 words, range = 4-15 words), but their length was matched between conditions (HC sentence frames: mean = 9.45 ± 2.20 words, range = 4-15 words; LC sentence frames: mean = 9.28 ± 2.03 words, range = 4-15 words; *p =* .455). The target stimuli were spoken by an artificial voice with a native Italian accent in one condition and an artificial voice with a foreign accent in the other condition. All stimuli of the native and foreign conditions were synthesized using the Microsoft Azure text-to-speech service. Two different speakers were used for the native and foreign-accented conditions. The prebuilt neural voice of an Italian speaker (Fabiola) was selected for the native-accented voice. The prebuilt neural voice of an Indian speaker (Neerja) was selected for the foreign-accent voice. For the foreign-accented voice, we used the artificial foreign accent used in Sala et al. (2024). The foreign accent was created by modifying three phonemes (/r/, /p/ and /k/) produced by the foreign-accented voice as /l/, /b/ and /ɢ/, respectively. The stimuli were synthesized starting from the IPA encoding of the words in Italian in order to minimize the impact of the phonetic variability between the native and the foreign voice. The phonological manipulation of the foreign-accented speaker was implemented by changing the target phonemes in the IPA encoding of the stimuli (e.g., the target stimulus was /kˈaldo/ for the native speaker voice and /g’aldo/ for the foreign speaker voice). The foreign-accented voice mispronounced the initial phoneme of all target stimuli. The mispronounced words did not correspond to any existing Italian word. The phonological manipulation was implemented to synthesize both the experimental stimuli and the familiarization speech of the foreign speaker. The spoken target stimuli had a similar duration between native and foreign accent conditions (native accent: 688 ±129 ms; foreign accent: mean = 700 ± 125 ms; *p =* .399). A more detailed description of the novel foreign-accent and of the stimuli generation procedure is reported in Sala et al. (2024).

Two face stimuli were selected from the AMFD (American Multiracial Face Database; Chen et al., 2021) to represent the native-accented speaker and the foreign-accented speaker. The AMFD includes face stimuli that have been rated by naïve observers on several dimensions, including “racial group” membership, evaluated on a 7-item Likert scale ranging from 1 (very atypical) to 7 (very typical). For the native accented speaker, we selected a face stimulus that we considered Italian-looking (White: mean = 6.22 ± 0.82) and evaluated as not typical in the other norms (Asian: mean = 1.65 ± 1.11; Black: mean = 1.68 ± 1.33; Latino/a: mean = 3.30 ± 1.73; Middle Eastern: mean = 2.57 ± 1.77; Biracial/Multiracial: mean = 3.62 ± 1.83). For the foreign-accented speaker, we selected a face stimulus evaluated as more typical for norms different from White/Caucasian (White: mean = 2.81 ± 1.73; Asian: mean = 2.54 ± 1.56; Black: mean = 2.56 ± 1.65; Latino/a: mean = 5.28 ± 1.19; Middle Eastern: mean = 4.40 ± 1.75; Biracial/Multiracial: mean = 4.60 ± 1.44). The two face stimuli had similar ratings in the other dimensions evaluated (see Table 1 and Table 2).

**Table 1.** AMFD ratings for the native- and foreign-accented speaker face stimuli.

|  | **Dominant** | **Trust** | **Smart** | **Warm** |
| --- | --- | --- | --- | --- |
| **Native speaker** | $2.97 \pm 1.85$ | $5.00 \pm 1.13$ | $5.24 \pm 1.09$ | $5.19 \pm 1.17$ |
| **Foreign speaker** | $2.79 \pm 1.48$ | $5.21 \pm 1.32$ | $5.09 \pm 1.11$ | $5.58 \pm 1.08$ |

**Table 2.** AMFD ratings for the native- and foreign-accented speaker face stimuli.

|  | **Attractive** | **Emotional**<br>**expression** | **Smile** | **Racial**<br>**ambiguity** |
| --- | --- | --- | --- | --- |
| **Native speaker** | $4.97 \pm 1.09$ | $5.78 \pm 0.82$ | $5.11 \pm 1.66$ | $2.86 \pm 1.62$ |
| **Foreign speaker** | $4.53 \pm 1.09$ | $5.82 \pm 0.93$ | $5.23 \pm 1.63$ | $3.89 \pm 1.37$ |

### Procedure and design

The experiment was carried out using PsychoPy (Peirce et al., 2019). Participants were seated in a comfortable chair in a soundproof room with a computer connected to a LCD monitor, speakers and a keyboard. During the familiarization phase, participants viewed a picture of the speaker’s face and listened to a one-minute speech in which the speaker introduced herself (see Sala et al., 2024). All participants were exposed to both speakers’ faces and listened to the corresponding accented speech. In the experimental phase, participants were instructed to silently read the sentence frames displayed on the screen. The speaker’s face was presented 2000 msec before sentence onset, 4.5 cm below the center of the screen and remained visible throughout the whole trial. In half of the trials, the face was replaced by a control stimulus, which was a scrambled version of the faces of the two speakers (see Fig. 1). Both the face and the control stimulus were 10 cm wide and 10 cm high. The presentation of each sentence frame started with a fixation cross appearing 4.5 cm above the center of the screen for 300 msec (450 ms before sentence onset). The sentence frames were presented word-by-word at a regular pace (300 msec word duration, 200 msec inter-word interval). We planned to present the target word 800 ms after the presentation of the last word of the sentence, however, due to a technical error this interval resulted to be 930 ms. A fixation cross followed the last word of the sentence frame. Participants were instructed to avoid blinks and eye movements during the experimental trials. Each trial was followed by a 1750 msec interval in which participants could blink. In the 25% of trials, a written question was presented to participants, asking them to judge whether they expected the word produced by the speaker or not, regardless of how it was pronounced. Participants were instructed to categorize the spoken targets as expected or not expected by pressing the ‘M’ or ‘C’ keys on the keyboard with their left and right index. In half of the questions, they were asked to respond to expected words with the index finger of their dominant hand. In the other half, they were asked to respond to unexpected words with the index finger of their dominant hand.

**Figure 1.**
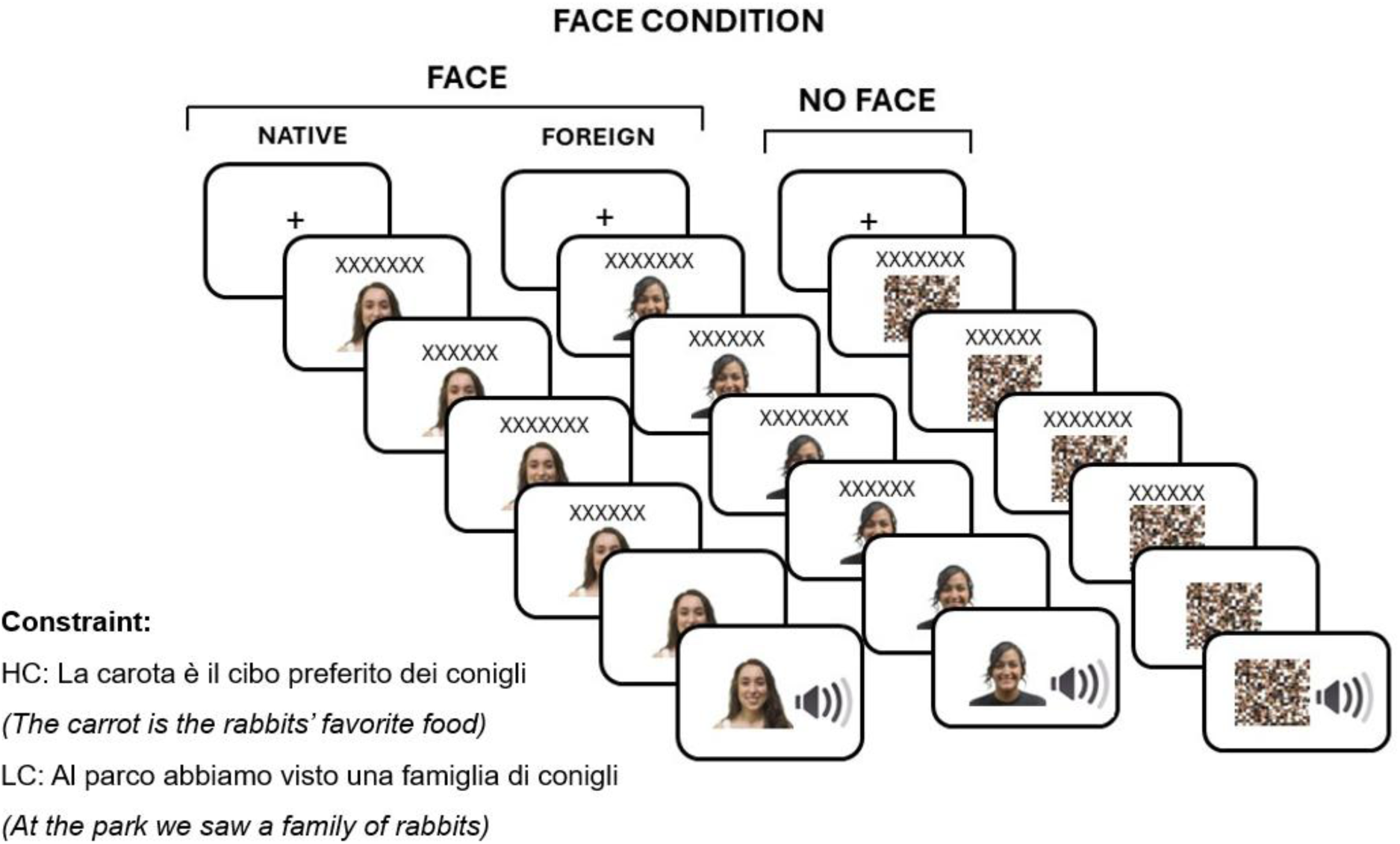
Schematic representation of the experimental paradigm and procedure. A trial consisted of a variable number of frames presented for 500 ms each (300 msec word duration, 200 msec inter-word interval). The number of frames depended on the length of each sentence. In each frame, a word was presented together with a visual stimulus that could be either the face of the native or foreign speaker or a control stimulus. The face cued the accent of the target word (in the example RANE/ frogs) that was presented auditorily, while the speaker’s face or the control stimulus remained visible. Sentences could be highly constraining (HC) or low constraining (LC) towards the target word. The face stimuli were selected from the American Multiracial Database (Chen et al., 2021) and are used under the CC-BY Attribution 4.0 International License.

Before starting the experimental session, participants completed 8 practice trials that were not part of the experimental materials. The whole experimental session lasted about 2 hours. Each participant was presented with 336 trials, and they had the possibility to take a break every 21 trials. Each target word was presented twice, once in the High and once in the Low constraining sentence frames. To avoid close repetitions, the experimental list was divided into two blocks, and each target word appeared only once per block. Within each block, 84 speech stimuli were spoken by the native speaker and 84 by the foreign speaker. The assignment of speech stimuli to the native or foreign-accented speaker was counterbalanced between blocks. The speaker’s accent was either cued or not cued by the speaker’s face, resulting in 42 trials per experimental condition. The order of the two blocks was counterbalanced between participants and the order of the trials within blocks was randomized. Since each target stimulus was presented in both native and foreign accents, and with or without the speaker’s face, four experimental lists were created, and the materials were rotated between conditions according to a Latin square design. Participants were randomly assigned to one of the lists.

### EEG data acquisition and preprocessing

Electroencephalogram was recorded with a system of 64 active Ag/AgCl electrodes (Brain Products), placed according to the 10–20 convention (ActiCap). Sixty of them were used as active electrodes (Fp1, Fp2, AF3, AF4, AF7, AF8, AFz, F1, F2, F3, F4, F5, F6, F7, F8, Fz, FT7, FT8, FC1, FC2, FC3, FC4, FC5, FC6, FCz, T7, T8, C1, C2, C3, C4, C5, C6, Cz, TP7, TP8, CP1, CP2, CP3, CP4, CP5, CP6, CPz, P1, P2, P3, P4, P5, P6, P7, P8, Pz, PO3, PO4, PO7, PO8, POz, O1, O2, Oz). The reference electrode was placed on the left earlobe. Three electrodes were used to record blinks and saccades (external ocular canthi and below the left eye). The setup was deemed acceptable if electrode impedance was below 20 kΩ at the end of electrode placement. The signal was amplified and digitized at a sampling rate of 500 Hz. Before the tasks, a resting state of 3 minutes was recorded. Descriptive statistics for the participants’ performance in the behavioral task were computed, to ensure that participants were indeed reading sentences and listening to words. One participant classified a high number of words (69%) in Low Constraint contexts as “predictable” and therefore was excluded. Pre-processing was performed using the MATLAB toolbox EEGLAB (Delorme & Makeig, 2004) and Fieldtrip (Oostenveld et al., 2011). EEG signals were offline re-referenced to the average activity of the right and left earlobes. A band-pass filter (0.5 – 80 Hz) was applied to the raw data. Noisy or flat channels were excluded (max 3 channels per participant). Segments with extreme muscle artifacts were also excluded. Subsequently, Independent Component Analysis (ICA) with dimensionality reduction to 60 components (PCA) was computed to detect and remove artifacts with known time-series and topographies (blink, saccades and power-line noise at 50 Hz). ICA was applied to band-pass filtered data (1-55 Hz) to optimize the identification of ocular artifacts and power-line noise. ICA weights were applied to the 0.5-80 Hz filtered data. Missing channels were interpolated (superfast spherical interpolation) after ICA correction. After a low-pass filter (30 Hz), 1200-ms epochs were extracted starting from 200 ms before the onset of the auditory stimuli. A pre-stimulus 200 ms baseline correction was applied to the extracted epochs. Trials including slow drifts, muscular activity or remaining ocular artifacts were manually removed (see Table 3). Two participants were excluded from the analyses due to the high number of rejected epochs (> 20%). Three more participants were excluded due to high residual alpha activity. The final sample included 42 participants (mean _age_ = 23.05 ± 2.97 years, 34 women).

**Table 3.** Accepted trials (mean and SD) for the different experimental conditions.

|  | High Constraint |  | Low Constraint |  |
| --- | --- | --- | --- | --- |
|  | Face | No Face | Face | No Face |
| **Native** | 41.02 ± 1.29 | 41.02 ± 1.26 | 41.19 ± 1.09 | 40.98 ± 1.32 |
| **Foreign** | 41.07 ± 1.47 | 40.83 ± 1.36 | 41.12 ± 1.40 | 41.02 ± 1.57 |

## Statistical analyses

The statistical analyses were performed using the statistical software R (R Core Team, 2023). The statistical analyses were performed in a priori determined N400 time-window (300-500 ms after word onset) in a cluster of centro-parietal electrodes (Cz, C3, C4, Pz, P3, P4) (Kutas & Federmeier, 2011; Nieuwland et al., 2018). The single-trial mean voltage in the N400 time-window was analyzed using linear mixed-effect models (Bates et al., 2015). We used a hierarchical model comparison approach to identify the best fitting model for our data (Heinze et al., 2018). The model comparison was based on the Akaike Information Criterion (AIC) and especially delta AIC and AIC weight as indexes of the goodness of fit. The AIC and AIC weight gives information on the models’ relative evidence (i.e., likelihood and parsimony), therefore the model with the lowest AIC and the highest AIC weight is to be preferred (Wagenmakers & Farrell, 2004). The model comparison included a base model with Constraint by Participant random slopes, and Participant and Item as random intercepts to account for participant-specific variability and item-specific variations (Baayen et al., 2008). The predictor’s order was established giving priority to the main effects over interactions and to the effects (i.e., Accent and Constraint) previously reported in the literature (Goslin et al., 2012; Grey & van Hell, 2017; Kutas & Federmeier, 2011;). The inclusion of predictors followed this order: (i) Accent (Native vs. Foreign); (ii) Constraint (HC vs. LC); (iii) Face (Face vs. No Face); (iv) Accent*Constraint; (v) Constraint*Face; (vi) Accent*Face; (vii) Accent*Constraint*Face. Main effects were estimated using sum coding (Brehm & Alday, 2022). The critical alpha was set to 0.05. Post hoc comparisons were performed using the contrast function of the emmeans package (Lenth et al., 2023). The p-values were adjusted using Bonferroni’s correction (Bonferroni, 1936).

## Results

### Behavioral results

In the 25% of trials, a written question was presented to participants, asking them to judge whether they expected the word produced by the speaker or not, regardless of how it was pronounced. The proportion of target words classified as “predictable” in control trials for the different experimental conditions is reported in Table 4.

**Table 4.**
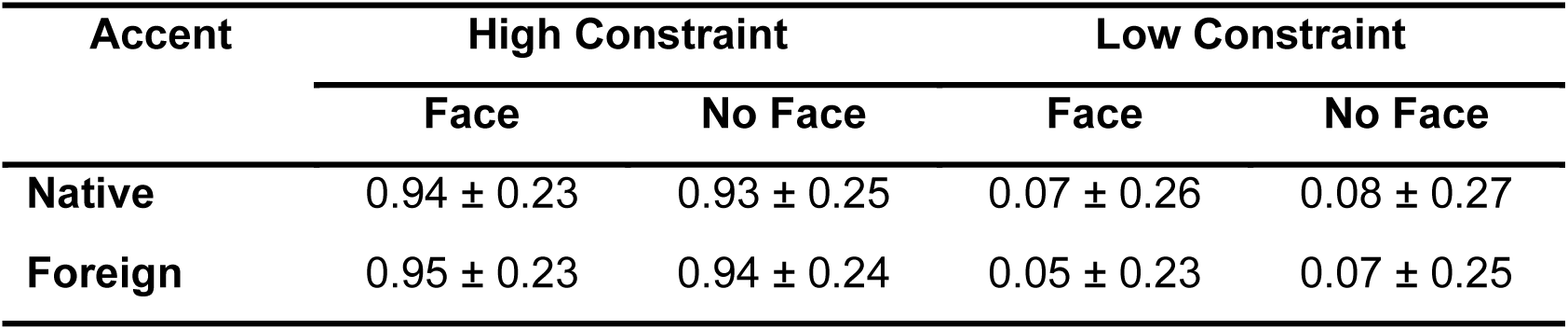
Proportion of target words classified as “predictable” in the different experimental conditions.

Participants seem to categorize most of the words in control trials with High Constraint contexts as *predictable*, and most of the words in control trials with Low Constraint contexts as *not predictable*, suggesting that they were paying attention to the stimuli. Moreover, this judgment doesn’t seem to be influenced by the presence of the face cue.

## ERP results

### Visual inspection of waveforms

Figure 2 shows the grand-average ERP waveforms in 9 representative electrodes (Fz, F3, F4, Cz, C3, C4, Pz, P3, P4) and the topographic maps of the experimental conditions.

**Figure 2.**
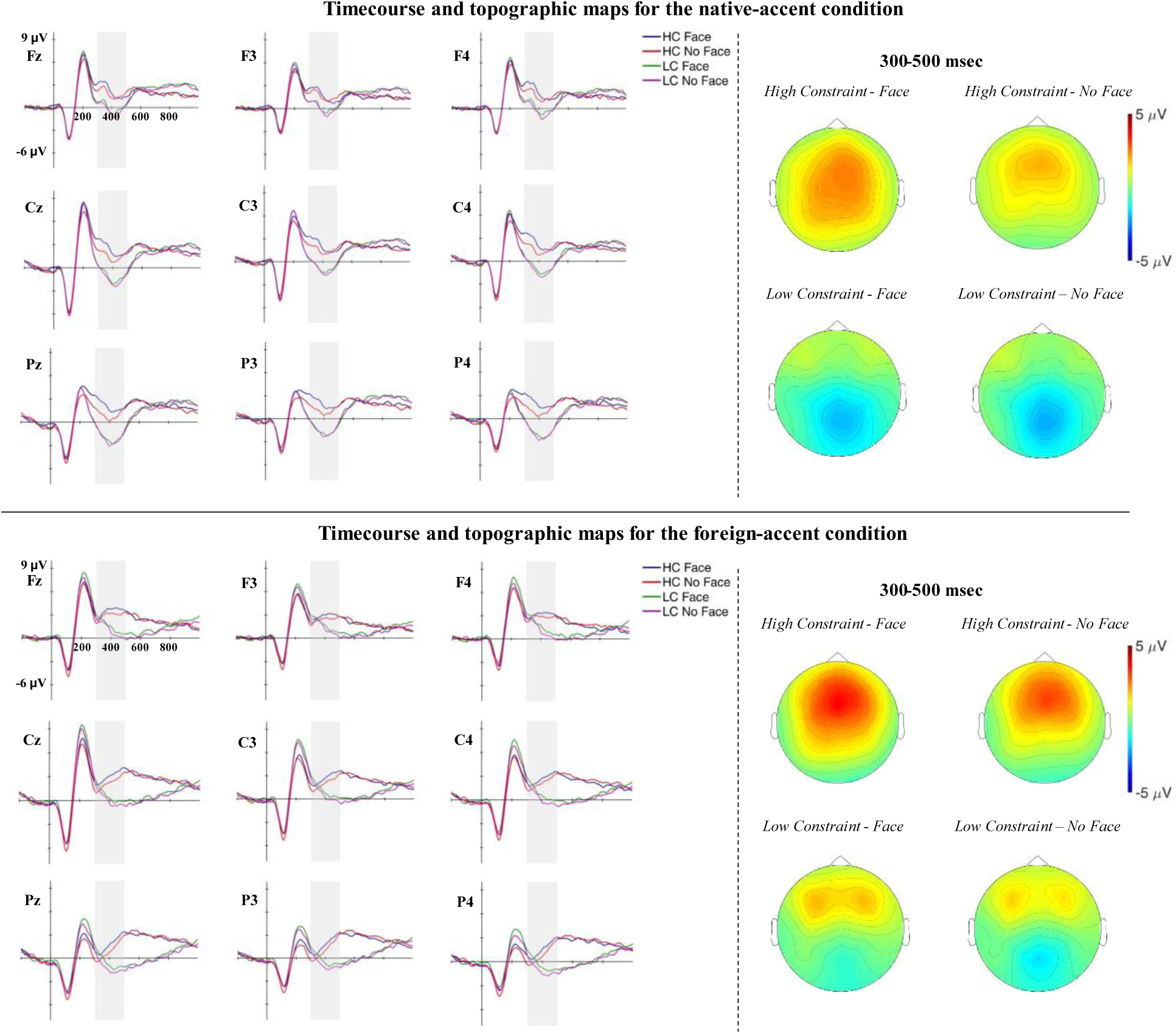
ERPs time-course and topographic maps for the native and the foreign-accented conditions. The grey pattern marks the N400 time-window (300-500 ms).

As predicted, we observed differences in the N400 time window according to the Constraint condition that are modulated by the presence of the face. However, the Constraint effect is present also in previous and successive time intervals and it seems to be different across the native vs. foreign accented condition (Figure 3).

**Figure 3.**
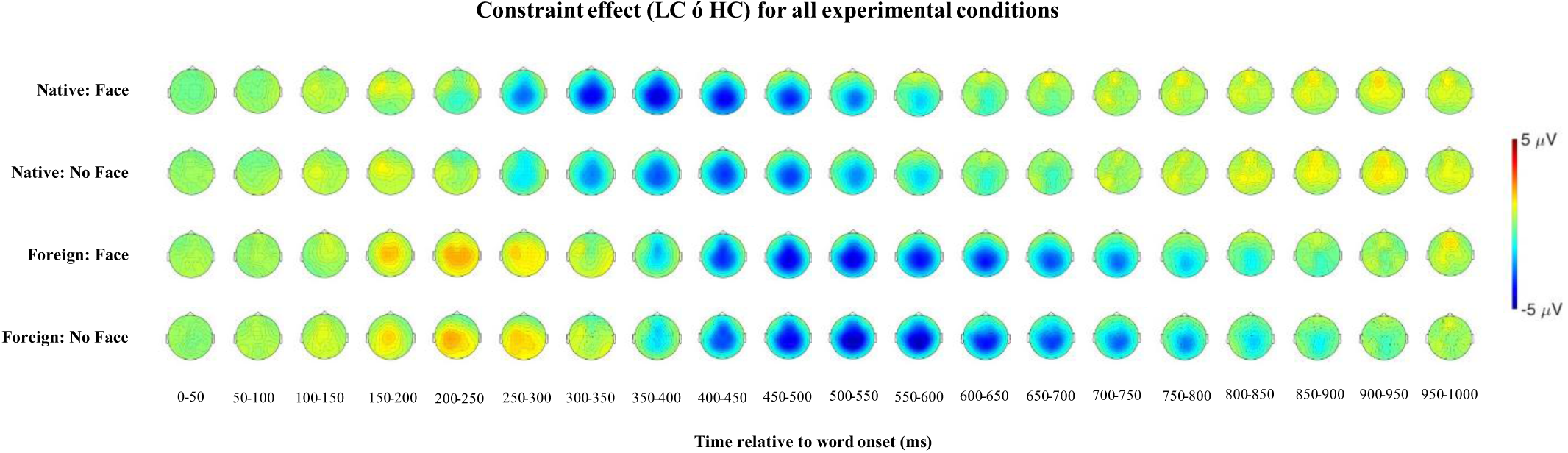
Topographic maps of the constraint effect according to Accent and Face conditions.

Compared to predictable words, unpredictable words in the native-accent condition present larger centro-parietal negativity starting from 250-300 ms. In the foreign-accent condition, unpredictable words present larger positivity between 150-300 ms, and larger centro-parietal negativity from 350-400 ms. Given this complex pattern we performed two types of analyses. A first a-priori planned analysis on the N400 time window and a second post-hoc temporal Exploratory Factor Analysis (EFA) to better characterize the ERP components underlying the effects observed after word onset. We computed separate EFAs for the native- and the foreign-accented conditions, as visual inspection of the grand-average waveforms points to noticeable differences in the ERP components elicited by these conditions.

**N400: 300-500 ms**

As reported in Table 5, the best-fitting model for N400 amplitude is Model 5:

Amplitude ∼ Accent + Constraint + Face + Constraint*Accent + Constraint*Face + (Constraint|Participant) + (1|Item)

**Table 5.** The comparison of LMER models predicting the N400 amplitude. Deviance = residual deviance; dAIC = difference between AIC of each model and the model with lower AIC; AICw = AIC weight.

| Models | Deviance | dAIC | AICw |
| --- | --- | --- | --- |
| M0. Amplitude ∼ (Constraint\|Participant) + (1\|Item) | 89293.77 | 166.67 | 0.0 |
| M1. Amplitude ∼ Accent + (Constraint\|Participant) + (1\|Item) | 89239.48 | 114.38 | 0.0 |
| M2. Amplitude ∼ Accent + Constraint + (Constraint\|Participant) + (1\|Item) | 89179.38 | 56.29 | 0.0 |
| M3. Amplitude ∼ Accent + Constraint + Face + (Constraint\|Participant) + (1\|Item) | 89150.66 | 29.56 | 0.0 |
| M4. Amplitude ~ Accent + Constraint + Face + Constraint*Accent + (Constraint\|Participant) + (1\|Item) | 89121.36 | 2.26 | 0.15 |
| **M5. Amplitude ~ Accent + Constraint + Face + Constraint\*Accent + Constraint\*Face + (Constraint\|Participant) + (1\|Item)** | **89117.09** | **0.0** | **0.46** |
| M6. Amplitude ~ Accent + Constraint + Face + Constraint*Accent + Constraint*Face + Accent*Face + (Constraint\|Participant) + (1\|Item) | 89116.87 | 1.78 | 0.19 |
| M7. Amplitude ~ Accent + Constraint + Face + Constraint*Accent + Constraint*Face + Accent*Face + Accent*Constraint*Face + (Constraint\|Participant) + (1\|Item) | 89114.73 | 1.63 | 0.20 |

The results of the best-fitting model (Model 5) are reported in Table 6. All predictors showing main effects are implied in interactions, therefore we focus on the latter. The interaction between Constraint*Accent indicates that the Constraint effect, namely smaller N400 amplitude for predictable words compared to unpredictable words, is larger for the native compared to the foreign-accented condition. Most importantly, the interaction between Constraint*Face indicates that cueing the speaker identity is associated with an increased Constraint effect (Fig. 4). Post hoc comparisons showed that cueing the speaker identity is associated with smaller N400 amplitude for predictable (b = - 0.770, z-ratio = -5.258, *p* < .001) but not for unpredictable words (b = - 0.343, z-ratio = -2.343, *p* = .076).

**Table 6.** Model results for the best-fitting model for the N400 amplitude.

|  | **Estimate** | **CI (95%)** | **Std. Error** | **t-value** | **p-value** |
| --- | --- | --- | --- | --- | --- |
| Intercept | 0.548 | [0.143 0.952] | 0.204 | 2.683 | 0.010 |
| Accent[Foreign] | 0.381 | [0.280 0.483] | 0.052 | 7.383 | < .001 |
| Constraint[HC] | 1.260 | [1.036 1.484] | 0.113 | 11.135 | < .001 |
| Face[Face] | 0.278 | [0.177 0.380] | 0.052 | 5.376 | < .001 |
| Accent[Foreign]*Constraint[HC] | -0.280 | [-0.381 -0.179] | 0.052 | -5.417 | < .001 |
| Constraint[HC]*Face[Face] | 0.107 | [0.005 0.208] | 0.052 | 2.064 | .039 |

**Figure 4.**
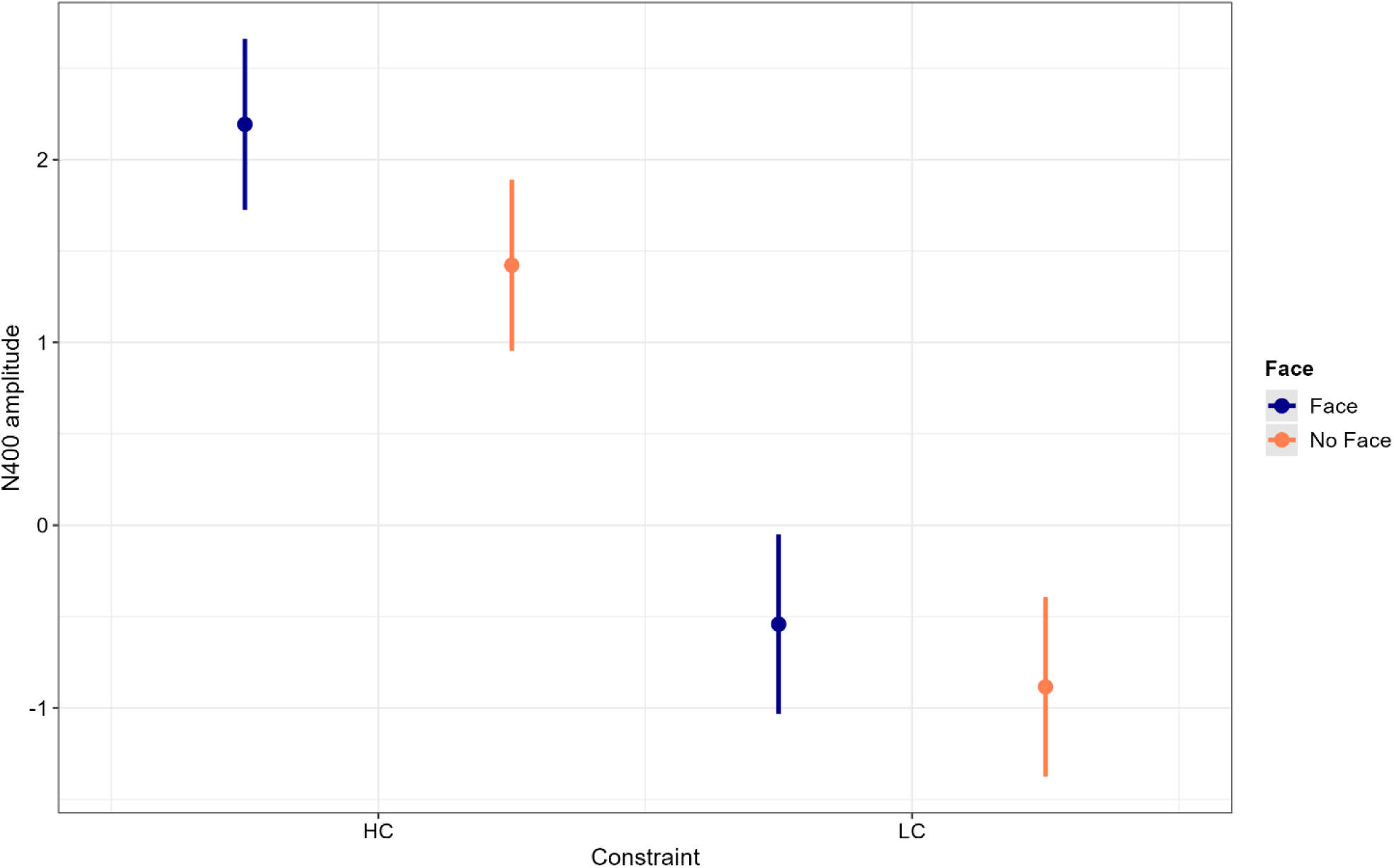
Model estimates for the interaction between Constraint*Face. The error bars indicate 95% confidence intervals.

### Interim discussion

In the present study, we aimed to investigate whether sentence contexts could be used to predict the phonological form of a highly predictable word. To address this, we capitalized on the fact that foreign-accented speakers often exhibit phonological errors to examine whether the prediction system can account for the phonological variability across speakers. In our experimental paradigm, participants read sentence frames in which the last word was produced by a native- or a foreign-accented speaker. The last word of the sentence could be predictable or not based on sentence context. Most importantly, during trial presentation speaker identity could be cued or not by an image of the speaker’s face, thus manipulating the availability of information about the phonological features of the upcoming target word before its presentation. We hypothesized that cueing speaker identity should be associated with a greater N400 predictability effect, reflecting easier processing of predictable words due to the pre-activation of the phonological word form. The findings from the main analysis supported our hypothesis: unpredictable words elicited a larger N400 compared to predictable words, and this N400 effect was larger when speaker identity was cued compared to when it was not. Crucially, post-hoc analyses showed that the face cue reduced the N400 amplitude for predictable words but not for unpredictable words. We found no evidence that the observed face cueing effect was modulated by the accent of the speaker (native vs. foreign). This result nicely aligns with previous behavioral data (Sala et al., 2024), further corroborating the involvement of phonological representations in linguistic prediction and suggesting that predictive processes are sensitive to interindividual differences in phonological features.

### Temporal Exploratory Factor Analysis

Figure 5 illustrates the grand-average ERP in Cz electrode for all the experimental conditions.

**Figure 5.**
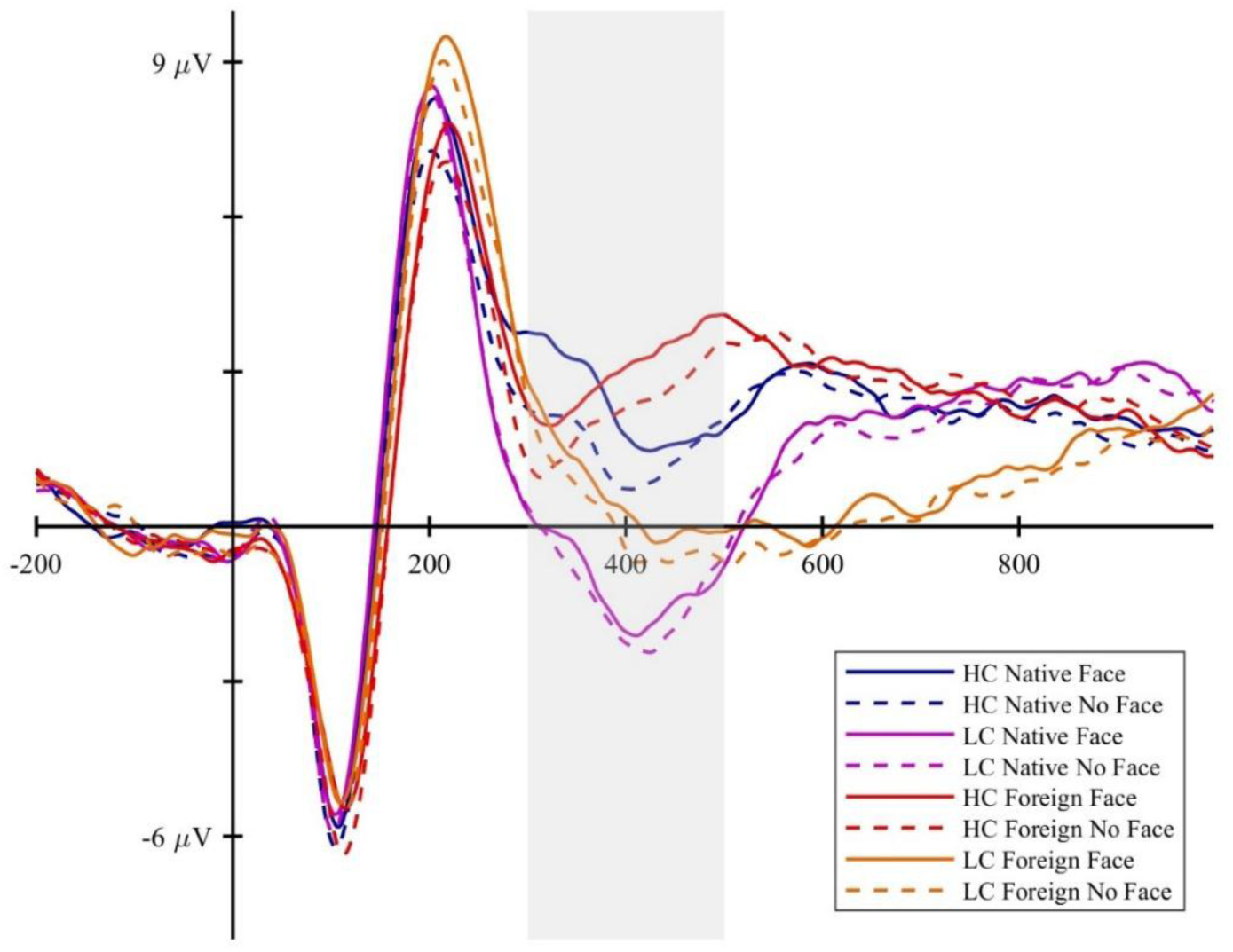
ERPs time-course in Cz electrode for all the experimental conditions, with the N400 time-window (300-500 ms) highlighted.

Visual inspection of the waveforms reveals a more complex pattern than hypothesized. First of all, starting from 300 ms after target onset, the non-subtracted waveforms seem to present a different shape for foreign-accented words compared to native-accented words. Moreover, the face cueing effect for predictable words seems to emerge prior to the N400 time-window, with positive deflections within the analyzed time-window.

The presence of positive deflections within the analyzed time-window may be due to the task used in the current study, in which participants were asked whether they expected the target word pronounced by the speaker in 25% of the trials. Brothers et al. (2015) used a task in which participants were asked to read medium-constraint sentences (cloze range 40–60%) that ended with either low cloze or medium cloze words (1 or 50% cloze probability, respectively). In all trials, participants were asked to indicate whether they expected the last word of the sentence. In their study, medium-cloze words were correctly predicted for 48.1% of passages (SD = 6.6%), while low-cloze words were consistently categorized as unpredictable. The ERP pattern showed that predicted medium-cloze targets compared to not-predicted medium-cloze targets were associated with a reduction of the N250 amplitude. The authors interpreted this effect as a facilitation of visual word form processing due to word form prediction. Alternatively, Nieuwland (2019) proposed to interpret this effect as a modulation of the P3b component, given that in the considered N250 time-window a strong positive deflection to predicted words was found. It is generally accepted that a distinction can be made between two subcomponents of the P300 response: the P3a (or novelty P3) and P3b (or target P3). The P3a is a positive deflection with a fronto-central distribution typically elicited in the oddball paradigm by infrequent non-target stimuli, while the P3b is characterized by a centro-parietal distribution and is elicited by task-relevant or target stimuli. In studies of language processing, the P3b has been observed in response to collocations (Molinaro & Carreiras, 2010), idioms (Vespignani et al., 2010), antinomies (Roehm et al., 2007), and expressions featuring predictable verbs in acceptability judgement task (Freunberger & Roehm, 2016). Its amplitude is thought to reflect the degree of correspondence between an internal representation and an external sensory stimulus during event categorization, with a larger amplitude reflecting a greater match, while its latency depends on the time required for stimulus evaluation and categorization (Kok, 2001; Kutas et al., 1977; McCarthy & Donchin, 1981). Predicted words in a prediction task could elicit a rapid and strong P3b response, and this may have also occurred in our experimental paradigm. When speaker identity was cued, predictable words may have elicited a greater P3b response reflecting a greater degree of correspondence between the internal representation of the upcoming word and the external input.

To better characterize the ERP components observed in our study, we conducted a Temporal Exploratory Factor Analysis (EFA). This approach was chosen because the traditional method of analyzing observed signal average across fixed time windows has limitations, as it assumes clear spatial and temporal separation between ERP components, which is not always the case (Luck, 2014). Temporal EFA is a statistical decomposition technique that can be used to identify and disentangle the constituent components of the observed ERPs (Dien, 2012; Dien & Frishkoff, 2005; Scharf et al., 2022)^1^. Temporal EFA allows the extraction of *factors* representing estimates of the underlying ERP *components,* based on statistical associations between sampling points. Factors are described by two coefficients: *factor loadings*, which represent the factor’s contribution to the voltage at a specific sampling point, and *factor scores*, which quantify the factor’s contribution to the observed ERP (for more details see Scharf et al., 2022).

Temporal EFA was computed following the procedure reported by Scharf et al. (2022). Average waveforms for each participant, electrode and condition were obtained by averaging the single trials.

The data were resampled at 250 Hz before running the Temporal EFA. Separate EFAs were computed for the native- and the foreign-accented conditions, as visual inspection of the grand-average waveforms points to noticeable differences in the ERP components elicited by these conditions. Conducting separate EFAs is preferable when *measurement invariance* cannot be assumed (Beauducel & Hilger, 2018; Meredith, 1993; Möcks, 1986), such as in cases of condition-related component structures or latency differences (Barry et al., 2016). Temporal EFA was computed on the trials-averaged waveforms, including the baseline pre-stimulus period and all the EEG channels (Dien, 2012). The EFA model was estimated using the covariance matrix of the sample points. The number of factors to be extracted was based on the Empirical Kaiser Criterion (EKC) (Braeken & van Assen, 2017; Y. Li et al., 2020). The rotated solution was estimated using Geomin rotation (Yates, 1987) with 30 random start values and the rotation parameter ɛ set to 0.01. We considered for the analysis only factors explaining more than 3% of the total variance.

We analyzed the *peak amplitude* (factor scores multiplied with the peak factor loading) of the factors reflecting ERP components of interest. For factors reflecting ERP components with a well-established topography, the cluster of electrodes analyzed was defined on a theoretical basis. We selected a cluster of centro-parietal electrodes (Cz, C3, C4, Pz, P3, P4) for factors reflecting components with a centro-parietal distribution (P3b and N400), while a cluster of fronto-central electrodes (Cz, C3, C4, Fz, F3, F4) was selected for factors reflecting components with a fronto-central distribution (N1, P2, P3a). We explored also factors reflecting slow waves extending beyond the N400 time-window. Given the heterogeneity of the scalp distribution of slow waves in the literature (see Van Petten & Luka, 2012), we selected a centro-parietal or a fronto-central electrode cluster based on the anterior or posterior distribution of the factor of interest. For each factor we run a repeated measures ANOVA, testing for following effects: i) Constraint, ii) Face, iii) Constraint*Face. Post-hoc comparisons were performed using the contrast function of the emmeans package (Lenth et al., 2023). P-values were adjusted using Bonferroni correction (Bonferroni, 1936).

### Native-accent Temporal EFA

In the native-accent condition, we extracted a 23-factor solution explaining 95% of the total variance. Fig. 6 illustrates the unstandardized factor loadings for the native-accent condition.

**Figure 6.**
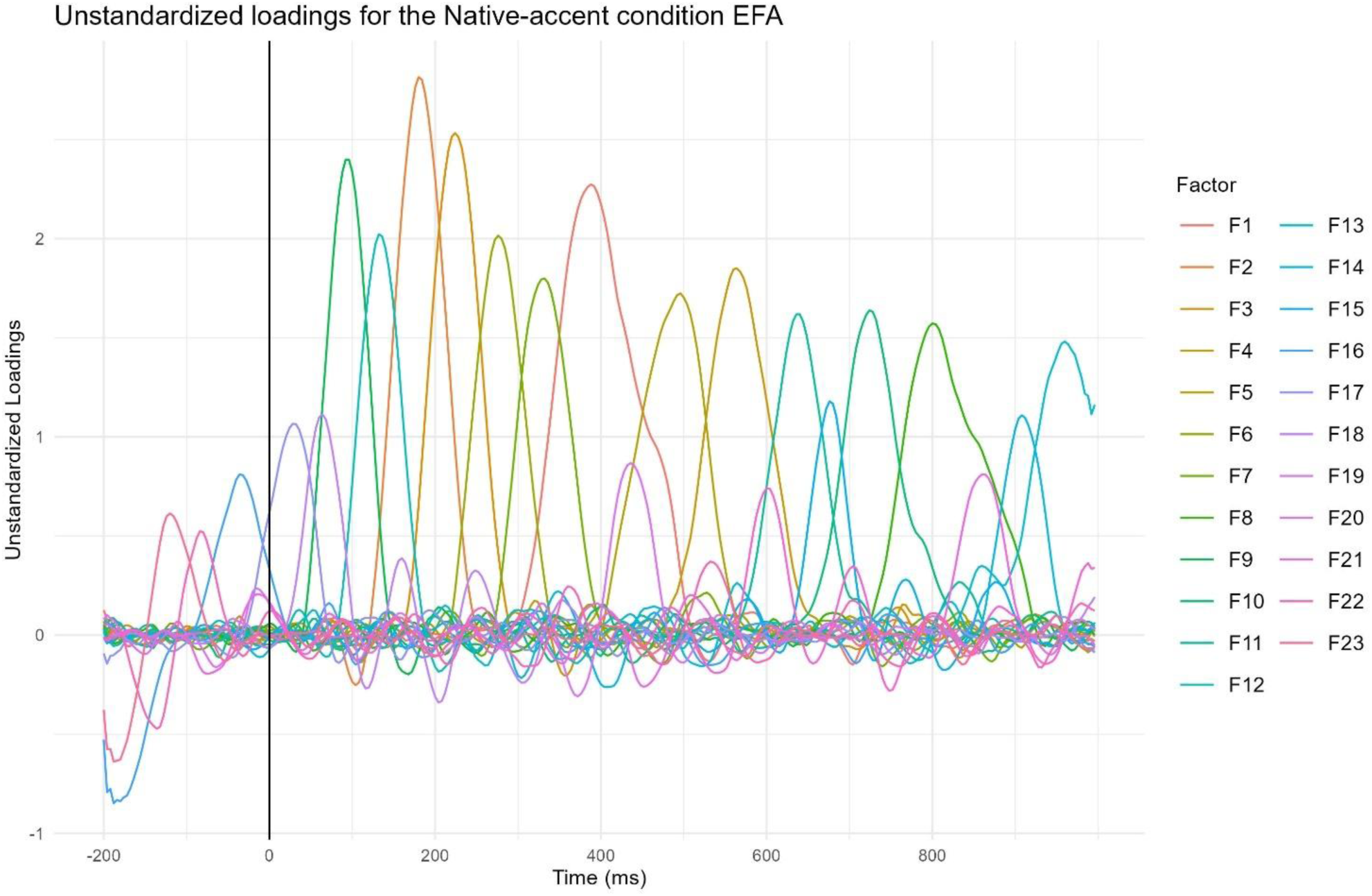
Unstandardized Factor Loadings after Geomin Rotation in the native-accent condition EFA. Each colored line represents the factor loadings of a factor. Higher factor loadings imply that the factor contributes more to the voltage at a sampling point. The factors are numbered by the amount of variance they explain.

The factors explaining more than 3% of the total variance are shown in Figure 7.

**Figure 7.**
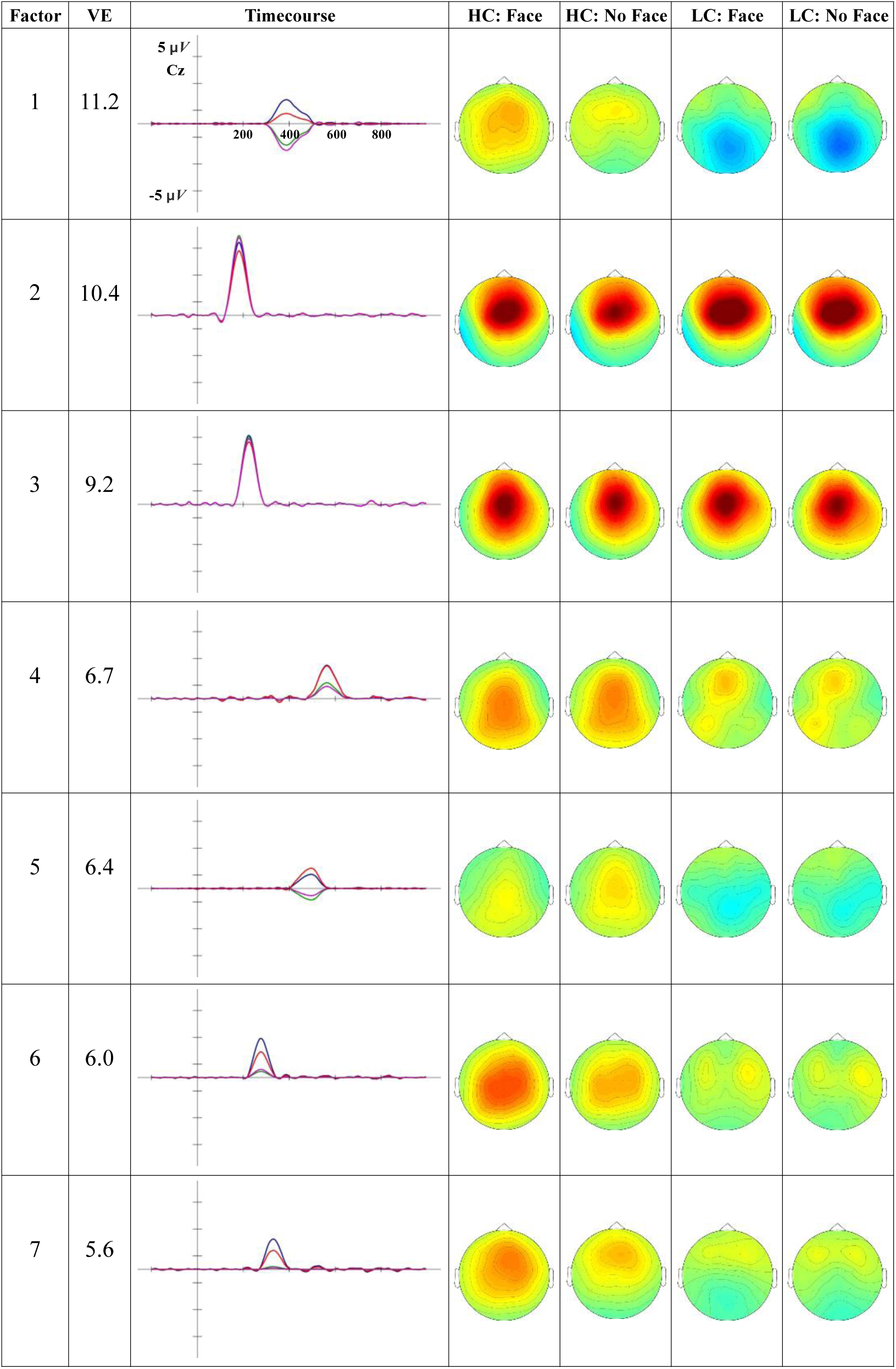
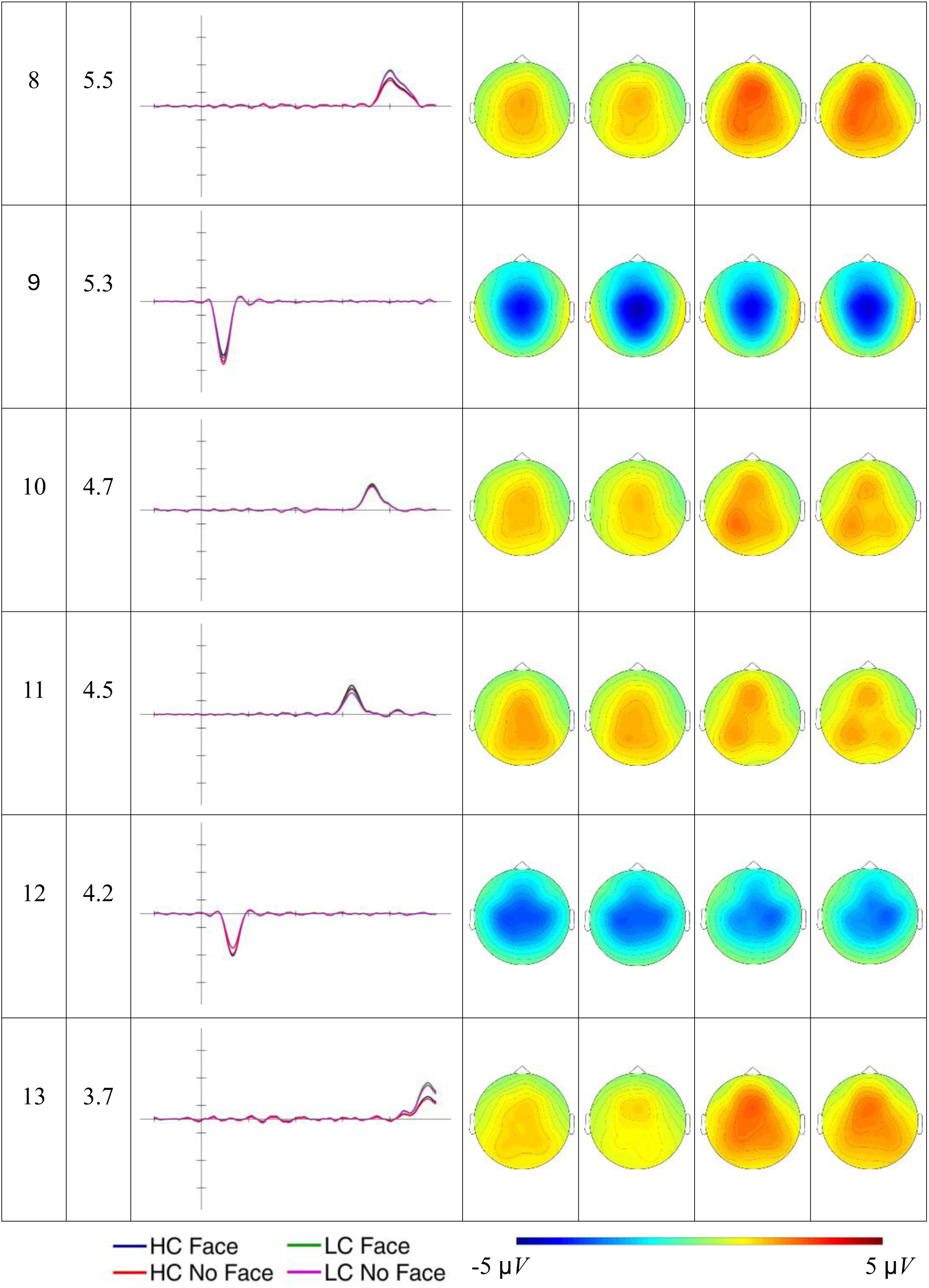
Timecourse and scalp distribution of the ERPs reconstructed factors in the different experimental conditions. VE: percentage of total variance explained by the extracted factor. The reconstructed ERP timecourse at the electrode Cz and the scalp distribution of the peak amplitude (factor scores multiplied with the peak factor loading) are reported.

Figure 8 presents the difference topographic maps for the Constraint effect in the factors explaining more than 3% of variance.

**Figure 8.**
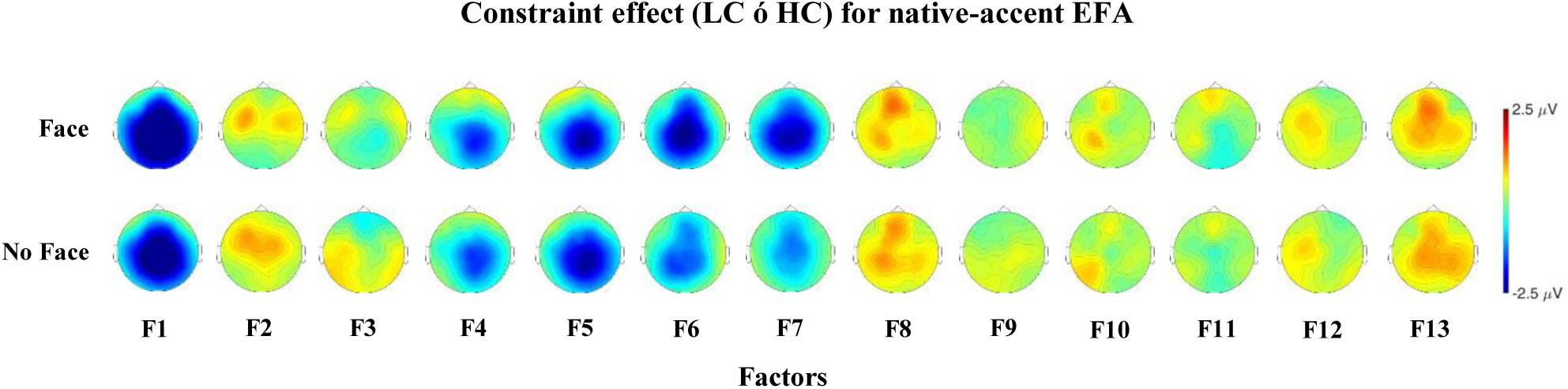
Topographic maps for the Constraint effect in the factors explaining more than 3% of variance.

Factors from 1 to 9 were considered reflecting ERPs components of interest and were analyzed. Table 7 reports the results of the models tested.

**Table 7.**
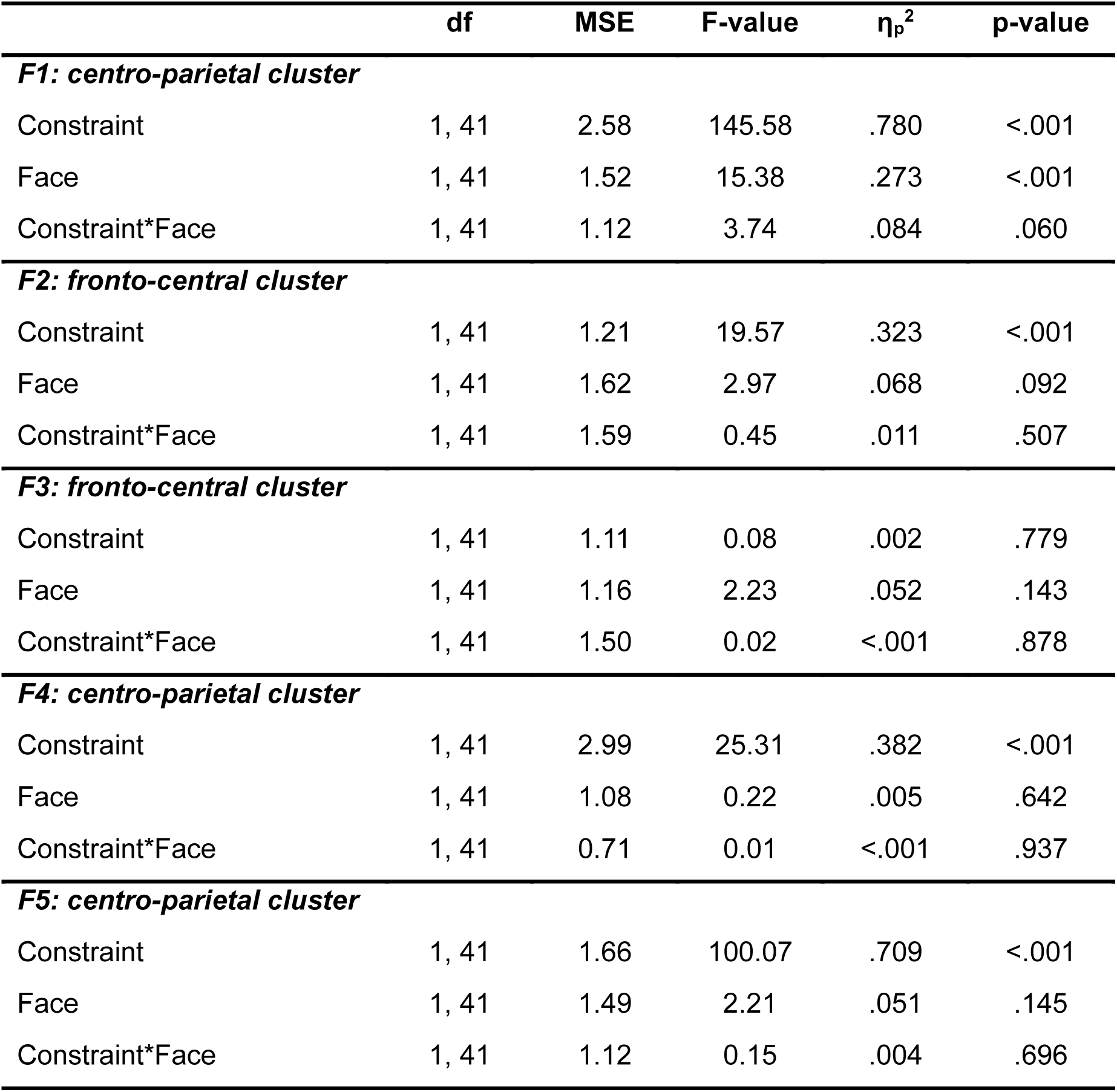
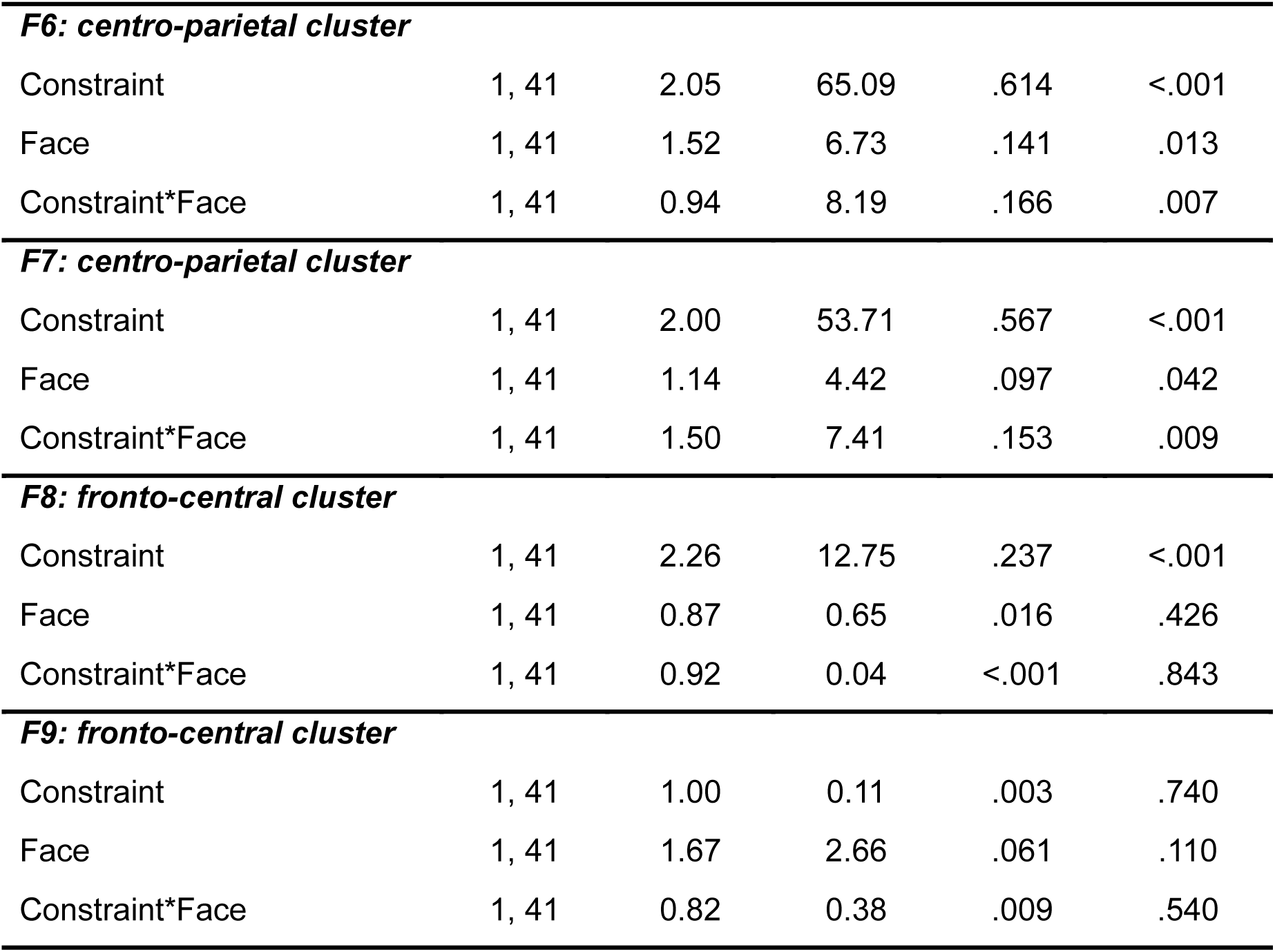
Results of the ANOVA models.

| | df | MSE | F-value | $\eta_p^2$ | p-value |
| --- | --- | --- | --- | --- | --- |
| ***F1: centro-parietal cluster*** |  |  |  |  |  |
| Constraint | 1, 41 | 2.58 | 145.58 | .780 | <.001 |
| Face | 1, 41 | 1.52 | 15.38 | .273 | <.001 |
| Constraint*Face | 1, 41 | 1.12 | 3.74 | .084 | .060 |
| ***F2: fronto-central cluster*** |  |  |  |  |  |
| Constraint | 1, 41 | 1.21 | 19.57 | .323 | <.001 |
| Face | 1, 41 | 1.62 | 2.97 | .068 | .092 |
| Constraint*Face | 1, 41 | 1.59 | 0.45 | .011 | .507 |
| ***F3: fronto-central cluster*** |  |  |  |  |  |
| Constraint | 1, 41 | 1.11 | 0.08 | .002 | .779 |
| Face | 1, 41 | 1.16 | 2.23 | .052 | .143 |
| Constraint*Face | 1, 41 | 1.50 | 0.02 | <.001 | .878 |
| ***F4: centro-parietal cluster*** |  |  |  |  |  |
| Constraint | 1, 41 | 2.99 | 25.31 | .382 | <.001 |
| Face | 1, 41 | 1.08 | 0.22 | .005 | .642 |
| Constraint*Face | 1, 41 | 0.71 | 0.01 | <.001 | .937 |
| ***F5: centro-parietal cluster*** |  |  |  |  |  |
| Constraint | 1, 41 | 1.66 | 100.07 | .709 | <.001 |
| Face | 1, 41 | 1.49 | 2.21 | .051 | .145 |
| Constraint*Face | 1, 41 | 1.12 | 0.15 | .004 | .696 |
| ***F6: centro-parietal cluster*** |  |  |  |  |  |
| Constraint | 1, 41 | 2.05 | 65.09 | .614 | <.001 |
| Face | 1, 41 | 1.52 | 6.73 | .141 | .013 |
| Constraint*Face | 1, 41 | 0.94 | 8.19 | .166 | .007 |
| ***F7: centro-parietal cluster*** |  |  |  |  |  |
| Constraint | 1, 41 | 2.00 | 53.71 | .567 | <.001 |
| Face | 1, 41 | 1.14 | 4.42 | .097 | .042 |
| Constraint*Face | 1, 41 | 1.50 | 7.41 | .153 | .009 |
| ***F8: fronto-central cluster*** |  |  |  |  |  |
| Constraint | 1, 41 | 2.26 | 12.75 | .237 | <.001 |
| Face | 1, 41 | 0.87 | 0.65 | .016 | .426 |
| Constraint*Face | 1, 41 | 0.92 | 0.04 | <.001 | .843 |
| ***F9: fronto-central cluster*** |  |  |  |  |  |
| Constraint | 1, 41 | 1.00 | 0.11 | .003 | .740 |
| Face | 1, 41 | 1.67 | 2.66 | .061 | .110 |
| Constraint*Face | 1, 41 | 0.82 | 0.38 | .009 | .540 |

F9 presents a negative deflection peaking at 92 ms with a central distribution, while F2 presents a positive deflection peaking at 180 ms with a fronto-central distribution. These factors seem to reflect the N1 and P2 components, respectively. While F9 was not modulated by any of the experimental conditions, F2 presents a Constraint effect, namely larger fronto-central positivity for unpredictable words compared to predictable words (b = -0.751, t-ratio_(41)_: -4.424, *p* < .001). F12 peaks at 132 ms and seem to reflect a sustained negativity for predictable words. F3 presents a positive deflection peaking at 224 ms with a fronto-central distribution. Its latency and topographical distribution closely resemble those of F2, suggesting that both factors may represent a temporal modulation of the P2 component, statistically divided into two factors. F3 was not modulated by any of the experimental conditions.

Two factors exhibit a positive deflection peaking around 300 ms: F6 and F7, which peak at 276 ms and 332 ms, respectively. F6 is more pronounced on the central electrodes, while F7 presents a fronto-central distribution. Both these factors seem to present a larger posterior positivity for predictable words compared to unpredictable words, that appear particularly pronounced when the face cue is present. This posterior positivity for predictable words may reflect a P3b response. The Constraint*Face interaction in F6 and F7 confirms that cueing the speaker identity is associated with a larger Constraint effect, namely larger positivity for predictable words compared to unpredictable words. Post-hoc comparisons showed that cueing the speaker’s face is associated with a larger centro-parietal positivity for predictable words (F6: b = -0.922, t-ratio_(41)_ = -3.747, *p =* 0.002; F7: b = -0.860, t-ratio_(41)_ = -3.372, *p =* .007) but not for not unpredictable words (F6: b = -0.064, t-ratio(41) = -0.270, *p >* .999; F7: b = 0.169, t-ratio(41) = 0.685, *p >* .999).

There are two factors that present a negative deflection for unpredictable words around 400 ms. F1 and F5 peak at 388 ms and 496 ms respectively and both seem to present a larger centro-parietal negativity for unpredictable words compared to predictable words, a pattern that is consistent with the N400 component. We observed a Constraint effect in both F1 and F5, confirming that unpredictable words elicit a larger centro-parietal negativity compared to predictable words. Moreover, F1 shows a Face effect (b = 0.747, t-ratio_(41)_: 3.922, *p* < .001), namely smaller negativity when the speaker identity is cued compared to when it is not cued. We also observed that both F1 and F5 present fronto-central positive activity for predictable words, alongside the posterior negative activity observed for unpredictable words. This pattern may reflect distinct ERP components with different spatial distributions, complicating the interpretation of the N400 amplitude modulations.

Among the factors emerging after the N400 time window (300–500 ms), F4 and F8 appear to be differently modulated by the predictability of the target word. F4 is associated with a larger centro-parietal negativity for unpredictable words compared to predictable words (b = 1.34, t-ratio_(41)_ = 5.031, *p* < .001), while F8 is associated with a larger fronto-central positivity for unpredictable words compared to predictable words (b = - 0.828, t-ratio_(41)_ = -3.571, *p* < .001). None of these factors seem to be modulated by the face cue.

### Foreign-accent Temporal EFA

In the foreign-accent condition, we extracted a 23-factor solution explaining 96% of the total variance. Fig. 9 illustrates the unstandardized factor loadings for the foreign-accent condition.

**Figure 9.**
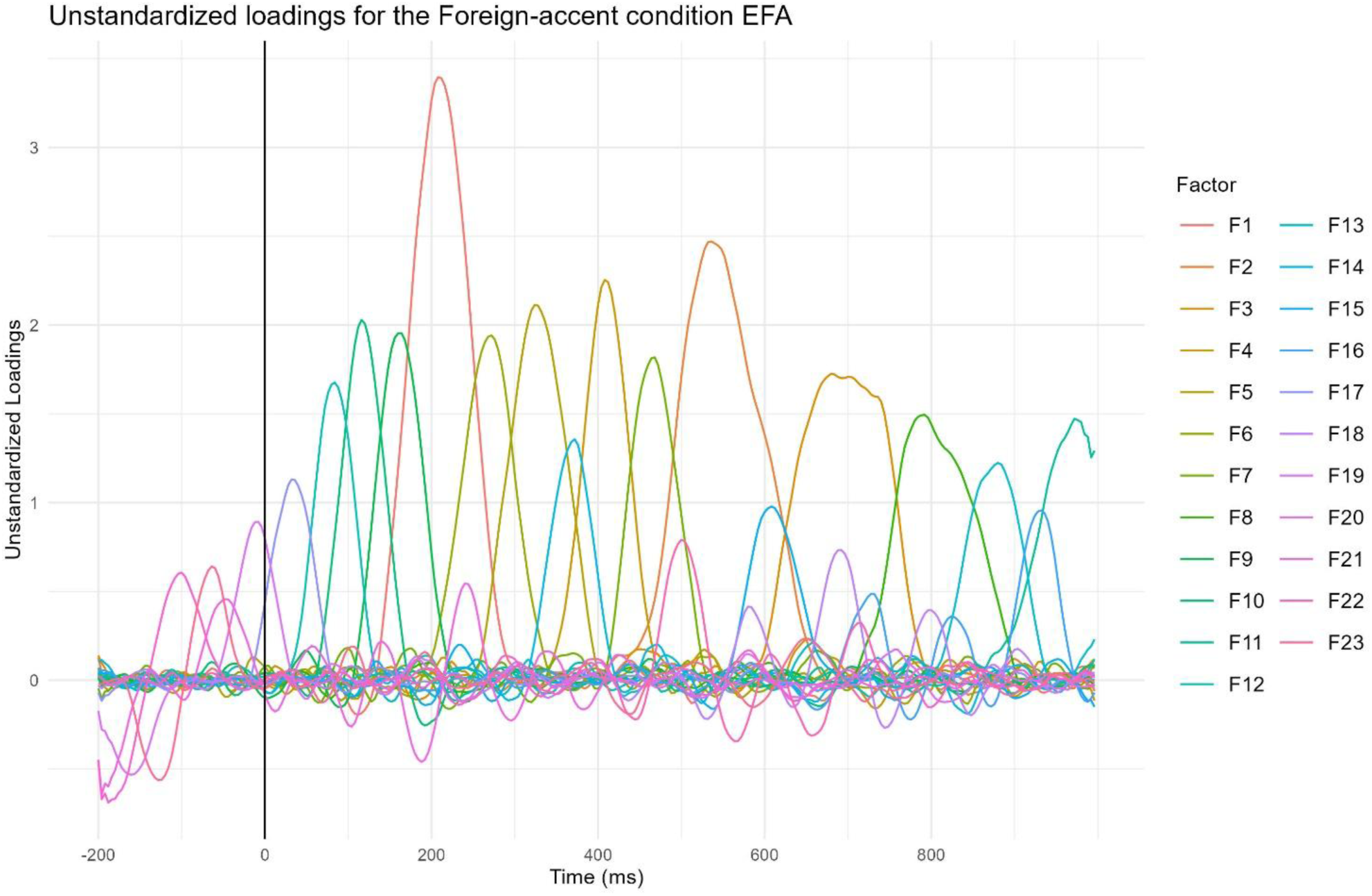
Unstandardized Factor Loadings after Geomin (0.01) Rotation in the foreign-accent condition EFA. Each colored line represents the factor loadings of a factor. Higher factor loadings imply that the factor contributes more to the voltage at a sampling point. The factors are numbered by the amount of variance they explain.

The factors explaining more than 3% of the total variance are shown in Figure 10.

**Figure 10.**
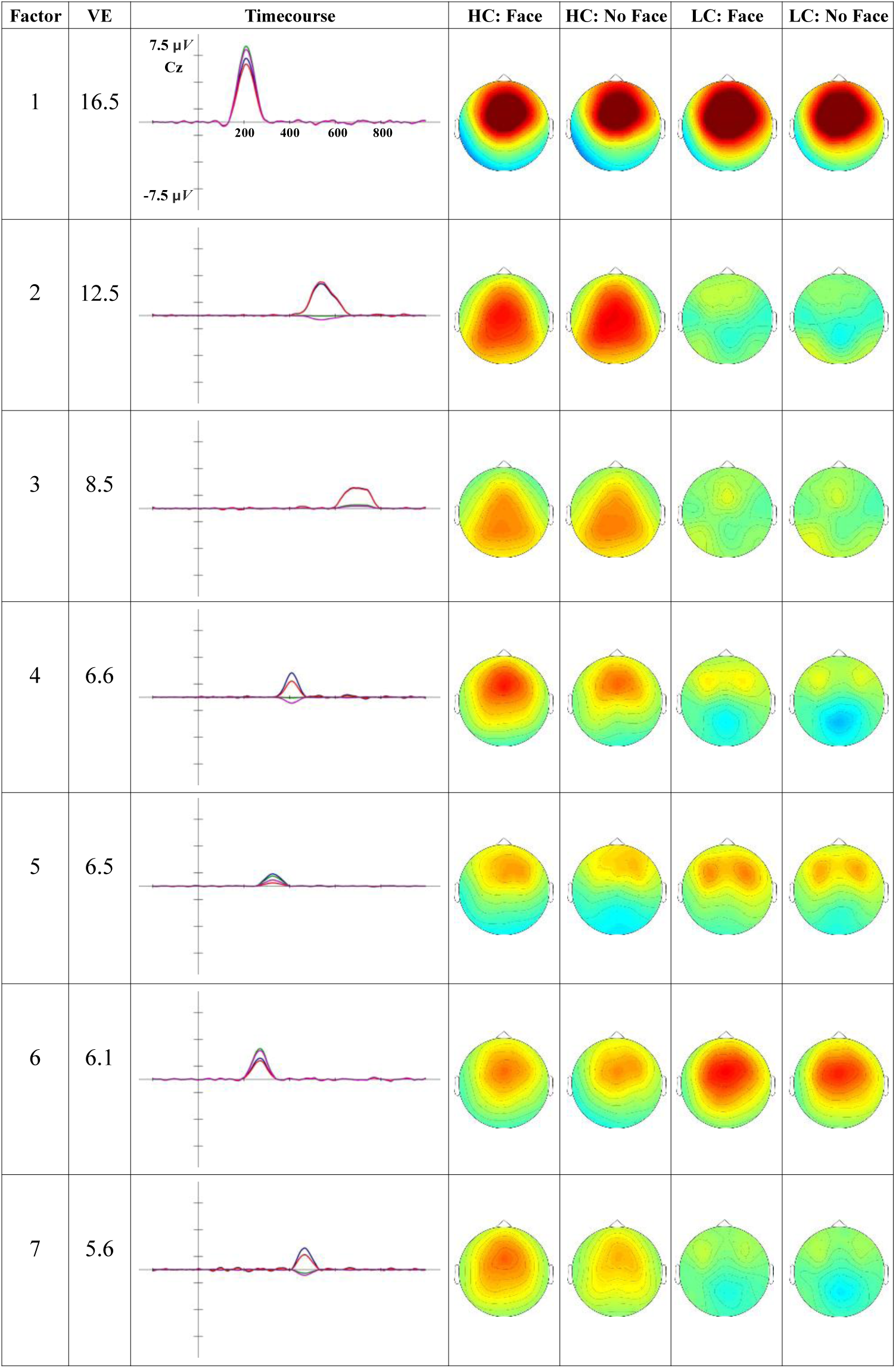
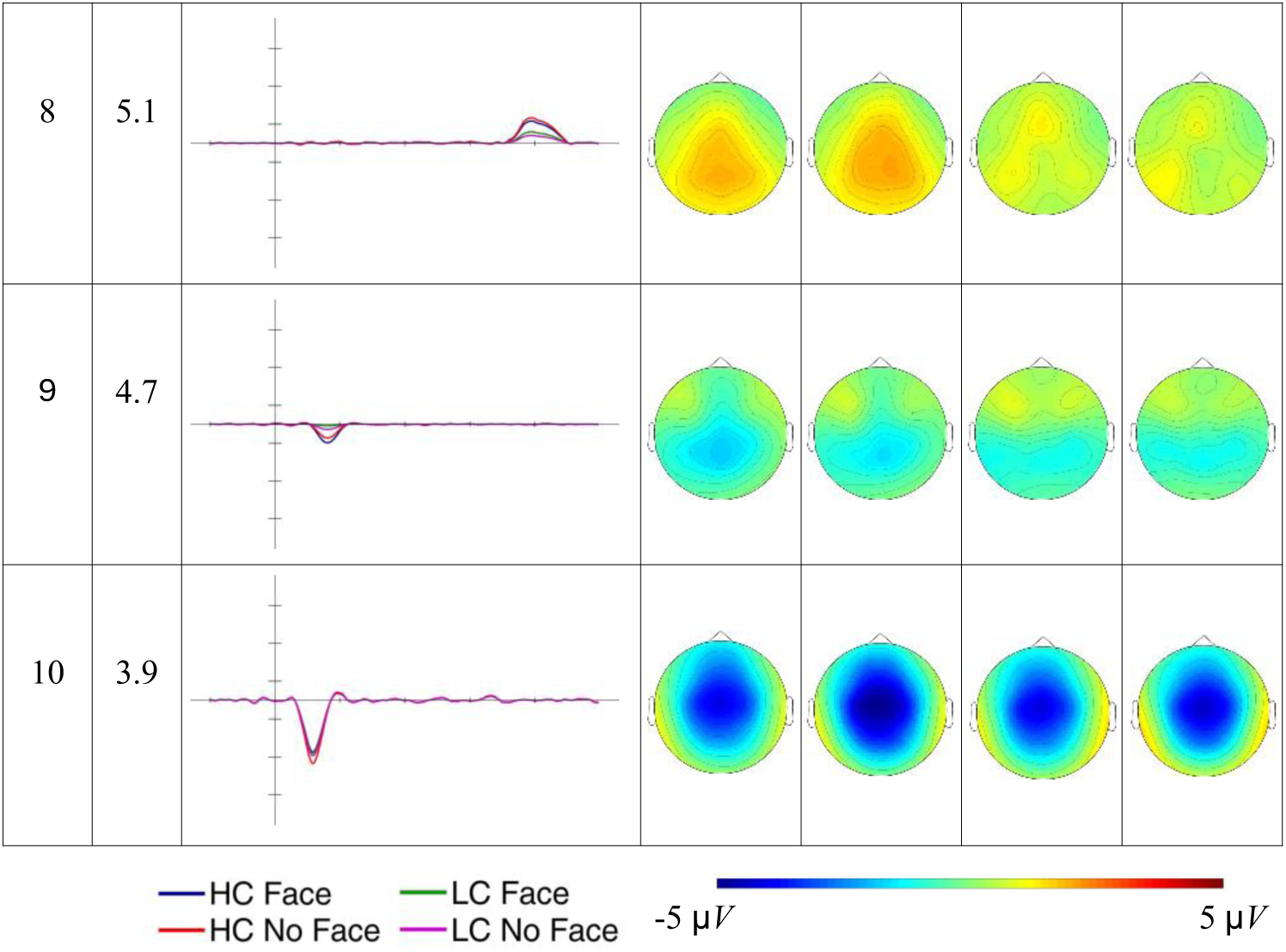
Timecourse and scalp distribution of the ERPs reconstructed factors in the different experimental conditions. VE: percentage of total variance explained by the extracted factor. The reconstructed ERP timecourse at the electrode Cz and the scalp distribution of the peak amplitude (factor scores multiplied with the peak factor loading) are reported.

Figure 11 presents the difference topographic maps for the Constraint effect in the factors explaining more than 3% of variance.

**Figure 11.**
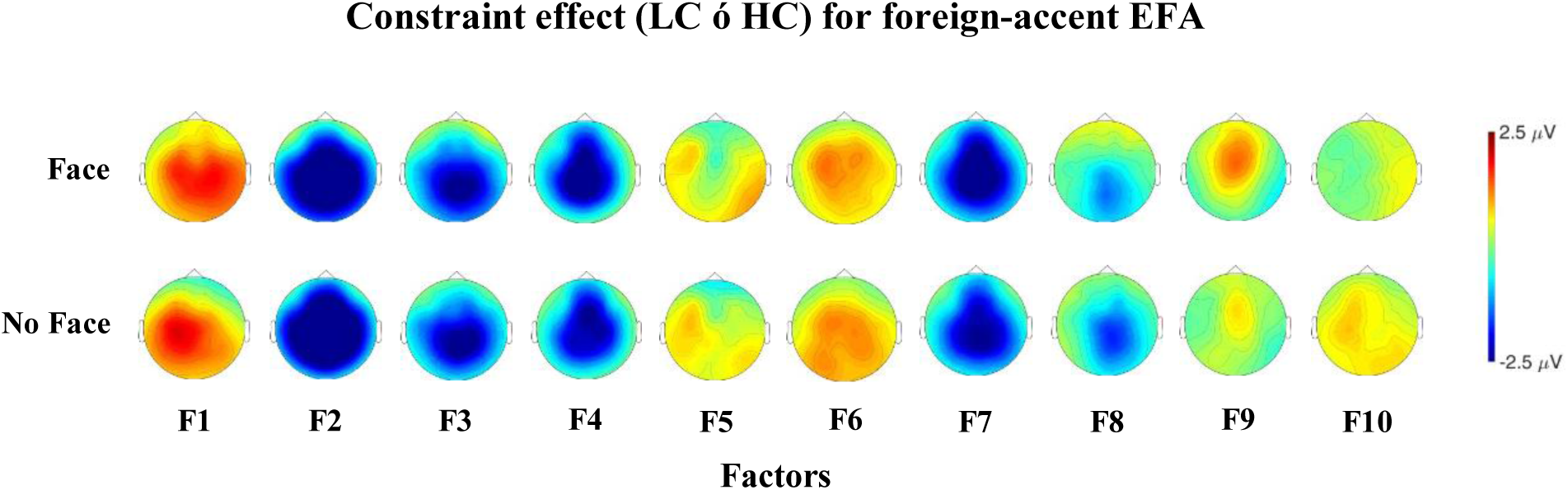
Topographic maps for the Constraint effect in the factors considered for the analysis.

Factors from 1 to 10 were considered reflecting ERPs components of interest and were analyzed. Table 8 reports the results of the models tested.

**Table 8.** Test statistics of the ANOVA models.

| | df | MSE | F-value | $\eta_p^2$ | p-value |
| --- | --- | --- | --- | --- | --- |
| **F1: fronto-central cluster** |  |  |  |  |  |
| Constraint | 1, 41 | 2.66 | 22.63 | .356 | <.001 |
| Face | 1, 41 | 1.51 | 3.35 | .076 | .074 |
| Constraint*Face | 1, 41 | 1.86 | 0.09 | .002 | .760 |
| **F2: centro-parietal cluster** |  |  |  |  |  |
| Constraint | 1, 41 | 4.35 | 148.95 | .784 | <.001 |
| Face | 1, 41 | 2.06 | 0.00 | <.001 | .991 |
| Constraint*Face | 1, 41 | 1.41 | 3.76 | .084 | .059 |
| ***F3: centro-parietal cluster*** |  |  |  |  |  |
| Constraint | 1, 41 | 2.26 | 91.20 | .690 | <.001 |
| Face | 1, 41 | 0.83 | 0.02 | <.001 | .901 |
| Constraint*Face | 1, 41 | 0.88 | 0.09 | .002 | .770 |
| ***F4: centro-parietal cluster*** |  |  |  |  |  |
| Constraint | 1, 41 | 4.06 | 56.96 | .581 | <.001 |
| Face | 1, 41 | 1.85 | 11.39 | .217 | .002 |
| Constraint*Face | 1, 41 | 0.82 | 0.88 | .021 | .353 |
| ***F5: fronto-central cluster*** |  |  |  |  |  |
| Constraint | 1, 41 | 2.51 | 0.71 | .017 | .405 |
| Face | 1, 41 | 1.16 | 9.18 | .183 | .004 |
| Constraint*Face | 1, 41 | 1.35 | 0.29 | .007 | .590 |
| ***F6: fronto-central cluster*** |  |  |  |  |  |
| Constraint | 1, 41 | 1.64 | 25.78 | .386 | <.001 |
| Face | 1, 41 | 0.89 | 3.18 | .072 | .082 |
| Constraint*Face | 1, 41 | 1.07 | 0.14 | .003 | .710 |
| ***F7: centro-parietal cluster*** |  |  |  |  |  |
| Constraint | 1, 41 | 2.61 | 93.03 | .694 | <.001 |
| Face | 1, 41 | 1.33 | 5.90 | .126 | .020 |
| Constraint*Face | 1, 41 | 1.33 | 0.69 | .016 | .413 |
| ***F8: centro-parietal cluster*** |  |  |  |  |  |
| Constraint | 1, 41 | 1.67 | 30.24 | .424 | <.001 |
| Face | 1, 41 | 1.02 | 0.01 | <.001 | .932 |
| Constraint*Face | 1, 41 | 1.40 | 1.59 | .037 | .214 |
| ***F9: fronto-central cluster*** |  |  |  |  |  |
| Constraint | 1, 41 | 1.36 | 16.97 | .293 | <.001 |
| Face | 1, 41 | 1.18 | 0.08 | .002 | .775 |
| Constraint*Face | 1, 41 | 1.45 | 2.64 | .060 | .112 |
| ***F10: fronto-central cluster*** |  |  |  |  |  |
| Constraint | 1, 41 | 1.24 | 3.76 | .084 | .059 |
| Face | 1, 41 | 1.39 | 5.84 | .125 | .020 |
| Constraint*Face | 1, 41 | 1.23 | 2.99 | .068 | .091 |

F10 presents a negative deflection peaking at 116 ms with a central distribution, while F1 presents a positive deflection peaking at 208 ms with a fronto-central distribution. These factors seem to reflect the N1 and P2 components, respectively. F10 exhibits a Face effect, showing reduced fronto-central negativity when the speaker’s identity is cued compared to when it is not (b = 0.439, t-ratio_(41)_: 2.416, *p* = .02). Similarly, F1 is influenced by Constraint, with a larger fronto-central positivity observed for unpredictable words compared to predictable words (b = -1.2, t-ratio_(41)_: -4.757, *p* < .001).

F9 peaks at 164 ms and appears to represent a prolonged frontal negativity for predictable words in contrast to unpredictable words (b = -0.742, t-ratio_(41)_: 4.12, *p* < .001).

Two factors exhibit a positive deflection peaking around 300 ms: F6 and F5, which peak at 272 ms and 324 ms, respectively. Both factors show a fronto-central scalp distribution, with F6 showing a more pronounced positivity for unpredictable words relative to predictable words. This pattern suggests that these factors may reflect a temporal modulation of the P3a response. We observed that F6 present a Constraint effect, with greater fronto-central positivity for unpredictable words relative to predictable ones (b = -1.0, t-ratio_(41)_: -5.078, *p* < .001). In contrast, F5 shows an effect of Face, displaying increased fronto-central positivity when the speaker’s identity is cued (b = -0.502, t-ratio_(41)_: 3.030, *p* = .004).

Two other factors, F4 and F7, exhibit a negative deflection for unpredictable words around 400 ms, peaking at 408 ms and 468 ms, respectively. Both factors seem to present a larger centro-parietal negativity for unpredictable words compared to predictable words, a pattern that is consistent with the N400 component. We observed a Constraint effect in both factors, confirming that unpredictable words elicit a larger centro-parietal negativity compared to predictable words (F4: b = 2.35, t-ratio_(41)_, 7.547, *p* < .001; F7: b = 2.40, t-ratio_(41)_ = 9.645, *p* < .001). Additionally, both are influenced by Face, as centro-parietal negativity is reduced when the speaker’s identity is cued (F4: b = 0.708, t-ratio_(41)_: 3.375, *p* = .002; F7: b = 0.432, t-ratio_(41)_: 2.429, *p* = .02). Moreover, in the foreign-accented condition, both candidate factors for the N400 component exhibit fronto-central positive activity for predictable words, alongside the posterior negative activity observed for unpredictable words. This pattern limits the possibility of drawing inferences about a potential modulation of the N400 amplitude.

F2, F3 and F8 emerge after the N400-timewindow (300-500 ms) and seem to reflect a stronger centro-parietal negativity for unpredictable words compared to predictable ones (F2: b = -3.93, t-ratio_(41)_: -12.205, *p* < .001; F3: b = 2.22, t-ratio_(41)_: 9.55, *p* < .001; F8: b = 1.1, t-ratio_(41)_: 5.499, *p* < .001).

### Summary of EFA analysis results

#### Native-accent condition

A main effect of Constraint was observed in factors we attributed to the P2, P3b, N400, and slow wave components. Specifically, the P2 was larger for unpredictable compared to predictable words. The P3b showed greater amplitude for predictable than for unpredictable words, while the N400 was smaller for predictable than for unpredictable words. Following the N400 time-window (300-500 ms), unpredictable words, compared to predictable words, elicited a sustained negativity, followed by a fronto-central positivity. A main effect of Face was found in a factor that may reflect the N400. The N400 was smaller when speaker identity was cued compared to when it was not cued. However, the presence of fronto-central positive activity for predictable words complicates the interpretation of these effects as a direct modulation of the N400 amplitude. Additionally, we observed a significant interaction between Constraint*Face in factors that may reflect the P3b. Cueing the speaker’s face was associated with a larger P3b response for predictable words but not for unpredictable words.

#### Foreign-accent condition

A main effect of Constraint was found in factors possibly reflecting the P2, P3a, N400 and slow waves. The P2 and the P3a were larger for unpredictable compared to predictable words, while the N400 was smaller for predictable than for unpredictable words. Following the N400 time-window (300-500 ms), unpredictable words elicited sustained negativity compared to predictable words. Unlike the native-accent condition, there was not a late fronto-central positivity for unpredictable words compared to predictable words. A main effect of Face was found in factors that may reflect the N1, P3a and N400. The N1 and the N400 were smaller and the P3a was larger when speaker identity was cued compared to when it was not cued. As in the native-accent condition, the presence of fronto-central positive activity for predictable words complicates the interpretation of potential modulations of the N400 amplitude. Unlike the native-accent condition, we did not observe any significant interaction between Constraint*Face in the factors analyzed.

## Discussion

In the current study, we hypothesized that cueing the speaker’s identity should facilitate the implementation of phonological predictions. Our main analysis supports this hypothesis, showing that cueing the speaker’s identity - hence the speaker-specific phonology of the upcoming word - is associated with a smaller N400 for predictable words but not for unpredictable words, and thus a larger predictability effect. However, this straightforward conclusion is complicated by the presence of positive deflections for predictable words within the N400 time-window. Previous research has shown that predicted words in a prediction task may elicit a P3b response (Brothers et al., 2015; Nieuwland, 2019). In the current study, in 25% of trials, participants were asked whether they expected the word pronounced by the speaker. Predictable words may have elicited a P3b response, which could have been stronger when speaker identity was cued, reflecting a greater match between the internal representation of the upcoming word and the external input. The results of the Temporal EFAs (Dien, 2012; Dien & Frishkoff, 2005; Scharf et al., 2022) provide evidence consistent with this hypothesis. In the native-accent condition, predictable words elicit a centro-parietal positivity compared to unpredictable words, possibly reflecting a P3b response. Moreover, this P3b response for predictable words was more pronounced when speaker identity was cued compared to when it was not. The selectivity of the face cue effect for predictable words suggests that the enhanced P3b response does not simply reflect the matching between the face cue and the speaker’s voice. As observed by Nieuwland (2019), it remains unclear whether the P3b response elicited by a task requiring an explicit decision on linguistic stimuli reflects recognition, word-form analysis or decision-related processes occurring after stimulus recognition. Our data provide some insights into this issue. In our task, participants categorized the final word of each sentence as predictable or not, regardless of how it was pronounced. While predictable words were inherently task-relevant, the presence of the face cue was not critical for deciding whether a stimulus was expected. Therefore, the observed modulation of the P3b in response to the face cue likely reflects facilitation in recognition or word-form analysis rather than a decision-making process. In the foreign-accented condition, the face cue effect appears as a modulation of the N1 and the P3a response. Cueing the speaker’s identity was associated with a suppression of the N1 response, regardless of sentence constraint. The N1 component is thought to reflect basic operations involved in constructing perceptual representations (Näätänen & Picton, 1987). It has been suggested that the N1 signals the detection of acoustic changes in the environment (Hyde, 1997), with its amplitude increasing in response to heightened attention to stimuli (Hillyard et al., 1973; Knight et al., 1981; Mangun, 1995; Ritter et al., 1988). Speech perception generally requires extracting acoustic cues from a continuous and transient signal and mapping them onto phonetic categories. This mapping is nondeterministic due to signal variability across contexts (c.f., lack of invariance) (Liberman et al., 1967). It has been proposed that speech perception relies on the listener’s knowledge of how linguistic units (words, syllables, phonetic categories, etc.) correspond to different distributions of acoustic cues (Kleinschmidt & Jaeger, 2015). From this perspective, speech perception is conceived as an inferential process under uncertainty, where the system generates probabilistic estimates of phonetic categories. Maintaining uncertainty about a category may be beneficial, as additional information becomes available as speech unfolds. Evidence suggests that later portions of a spoken stimulus can influence the interpretation of earlier portions (cf. right-context effects in word recognition; Bard et al., 1988; Connine et al., 1991; Dahan, 2010; Grosjean, 1985), suggesting that listeners retain some degree of uncertainty about speech signal for a limited time. The suppression of the N1 observed for foreign-accented words may reflect this process, wherein the system delays categorization to wait for additional information that facilitates mapping the speech signal onto linguistic categories. Listeners may allocate less attentional resources at the first phoneme of the target word, where foreign-accented speech diverge from the standard phonology, awaiting further information that could guide interpretation. Additionally, foreign-accented words seem to elicit a P3a rather than a P3b response. The P3a has been linked to processes related to the orienting response (Cycowicz & Friedman, 1997; Friedman et al., 1998; Knight & Nakada, 1998). These processes do not simply reflect the detection of a deviant or novel event but occur after the brain has identified the deviation, bringing it to conscious awareness for further evaluation and determining an appropriate response. The P3a was likely elicited in the foreign-accent condition due to participants’ limited familiarity with foreign-accented speech. The P3a exhibited an earlier modulation due to target word predictability, with unpredictable words eliciting a greater response than predictable words. It also showed a later modulation related to the face cue, with a greater response when speaker identity was cued compared to when it was not. The earlier modulation of the P3a may reflect an increased resource allocation for unpredictable words compared to predictable words. The later modulation of the P3a may be associated with a delay in the initial categorization of the speech signal, with the system allocating more resources to extract information that facilitates phonetic categorization at word onset. Extending beyond the canonical N400 time-window, in both accent conditions, we observed a sustained centro-parietal negativity for unpredictable words compared to predictable words. In language processing tasks, slow negative waves have been observed at the end of sentences containing an earlier ungrammaticality that elicits a P600 (De Vincenzi et al., 2003; Molinaro et al., 2008; Osterhout & Holcomb, 1992; Osterhout & Mobley, 1995; Osterhout & Nicol, 1999). Stowe et al. (2018) proposed that sentence-final negativities can be interpreted as the end of a slow wave elicited by the original ungrammaticality rather than a response linked to the processing of the final word per se. Sentence-final negativities emerge primarily when a decision about the linguistic stimuli is required, suggesting that they may be related to maintaining information relevant to the decision task. This aligns with ERP studies showing slow negative waves during tasks that involved activation of working memory (Lang et al., 1987; McCallum et al., 1988; Peronnet & Farah, 1989; Ruchkin et al., 1988, 1995). In low constraining sentence contexts, where predictions about upcoming words are weaker or more uncertain, the system may maintain multiple potential candidates in working memory. As a result, the decision-making process for the target word becomes more complex compared to highly constraining contexts, where one lexical candidate is far more likely than any alternative, leading to a prolonged negativity for unpredictable words. Finally, in the native-accented condition, we observed a late fronto-central positivity for unpredictable words compared to predictable words, emerging around 700 ms after word onset. Late frontal positivities can be elicited by semantically congruent sentence completions and are usually more pronounced for unpredictable words than for predictable words (Van Petten & Luka, 2012). It has been proposed that late frontal positivities reflect disconfirmed lexical predictions rather than general conceptual predictions (Thornhill & Van Petten, 2012; Van Petten & Luka, 2012). In our study, we observed a late frontal positivity for native-accented words, whereas no such positivity was observed for foreign-accented words. In the native-accent condition, predictable words likely matched the expected lexical form, whereas unpredictable words did not. For foreign-accented words, their non-standard phonology may have hindered lexical access, which, in turn, made the comparison between expected lexical form and pronounced word more challenging.

### What do our data say about linguistic prediction models?

The results of our study provide compelling evidence supporting the involvement of phonological representations in prediction, at least in highly informative and constraining contexts. The analysis of the observed amplitude in the N400 time-window suggests that cueing speaker identity is associated with an easier processing of predictable words due to the pre-activation of the phonological word form. Temporal EFA revealed substantial differences between the native and the foreign-accented conditions in the processes by which comprehenders use available information to predict upcoming speech. In the native accent condition, the face cueing effect for predictable words appears to be driven by a larger P3b response, reflecting a greater degree of correspondence between the internal representation of the upcoming word and the external input. In the foreign-accented condition, we did not find a clear factor associated with a face cueing effect for predictable words. Instead, we observed that cueing speaker identity is associated with a smaller N1 and a larger P3a, suggesting that comprehenders, when expecting non-standard pronunciations, adopt a more flexible processing strategy, maintaining uncertainty about the speech signal while leveraging contextual information to facilitate comprehension. Our findings help elucidate the mechanisms underlying phonological predictions. A standard assumption in the psycholinguistic literature is that activation of a lexical item spreads to similar or related items, due either to learned association in semantic networks (Anderson, 1983; Collins & Loftus, 1975; Hutchison, 2003) or overlapping semantic features between concepts (McRae et al., 1997). More recent cognitive frameworks postulated the presence of recurrent interactions between different levels of representation. As a result, the activation of a semantic representation could percolate top-down influencing the processing of the phonologic (Huettig et al., 2022) or orthographic (Kim & Lai, 2012; Molinaro et al., 2013) word features. These accounts extend beyond word-to-word semantic priming to encompass a broader sense of priming, which includes facilitation based on both linguistic and non-linguistic contextual information, considering the pre-activation of low-level features as a downstream consequence of the (pre-)activation of lexical-semantic representations. In our experimental paradigm, the speaker’s face serves as cue to the phonological properties of the upcoming word before its presentation. Although spreading of activation accounts acknowledge that non-linguistic context can influence phonological predictions by making the pre-activation of lexical-semantic representations more or less focal, they fail to explain how a phonologically informative non-linguistic cue can modulate the pre-activation of phonological representations. Prediction-by-production accounts represent a valuable framework for understanding how listeners use the available information to generate phonological predictions. According to these models, prediction during comprehension relies not only on associative mechanisms but also on processes traditionally attributed to language production (Huettig, 2015; Pickering & Gambi, 2018; Pickering & Garrod, 2007, 2013). Neurophysiology studies comparing the processing of high- and low-constraining sentence frames followed by a picture to be produced or by a word to be perceived, showed very similar brain time-frequency modulations between planning in word production and predicting during comprehension (Gastaldon et al., 2020), and that such neural markers are altered when speech planning is impaired (Gastaldon et al., 2023). These findings suggest that the language production network contributes to some extent to the prediction of the last word in high constraint sentences. Although prediction-by-production accounts are not entirely in agreement regarding the processes and representations involved in linguistic prediction (for a review, see Gastaldon et al., 2024), they often assume a role of event simulation in the generation of predictions. Pickering & Garrod (2013) proposed that listeners covertly imitate the unfolding utterance in order to derive an internal representation of the speaker’s percept, which constrains the prediction of upcoming speech. In this account, predictions are generated through forward models of the upcoming speech in the form of “impoverished” production representations. Later accounts have diverged from Pickering & Garrod (2013), arguing that prediction relies on fully-fledged production representations (Huettig, 2015; Pickering & Gambi, 2018) and that event simulation is a separate mechanism interacting with production (Huettig, 2015). Despite the different status of event simulation in prediction-by-production accounts, this mechanism could help to explain how listeners use the available information to generate phonological predictions, especially for the native accented speaker. The accuracy of an internal model of the speaker should increase with the comprehender’s similarity to the speaker (Pickering & Garrod, 2013). The internal model may incorporate different types of speaker-related information, such as shared background knowledge (Pickering & Gambi, 2018) and phonological features, allowing to predict how the speaker will pronounce the upcoming word. In our paradigm, comprehenders likely relied more on simulation with the native-accented speaker, as they could construct a more accurate internal model of the speaker. The literature includes other prediction models that could account for the implementation of phonological predictions without relying on event simulation or word production mechanisms. Kuperberg and Jaeger introduced a multi-representational hierarchical generative model (Kuperberg, 2016; Kuperberg & Jaeger, 2016) in which comprehenders rely upon internal generative models – defined as a set of hierarchically organized internal representations – to probabilistically pre-activate information at multiple levels of representation. The pre-activation of linguistic representations maximizes the probability of accurately recognizing the incoming information. Internal representations are derived from a limited number of observations, encompassing both linguistic and non-linguistic information, and they may also include knowledge of the speaker’s sound structure (Connine et al., 1991; Szostak & Pitt, 2013).

Listeners may learn different generative models corresponding to different statistical environments (Kleinschmidt & Jaeger, 2015), and these models will likely be more precise with increasing familiarity with the speaker.

In a previous behavioral study (Sala et al., 2024), we used a similar experimental design to investigate whether speaker identity (native vs. foreign) is used to implement phonological predictions. Participants were required to perform a lexical decision task on the spoken target word. In the case of foreign accent, participants were explicitly asked to accept mispronounced words as real. The results showed that cueing the speaker’s identity speeded up the RTs for predictable words, regardless of the speaker’s accent, suggesting that phonological predictions were similarly instantiated for both the native and the foreign-accented speaker. In the present study, by tapping onto electrophysiological responses, we found clearer evidence of phonological predictions for the native-accented speaker. This apparent discrepancy in the results can be explained by the different tasks used. In the behavioral study, participants were required to discriminate between real words mispronounced by the foreign-accented speaker and non-words, whereas in the present study, participants were asked to indicate whether they expected the spoken word regardless of its pronunciation. Participants may have implemented phonological predictions more flexibly in the lexical decision task, as such predictions could be useful in distinguishing between mispronounced real words and non-words pronounced by the foreign-accented speaker. This aligns with the Kuperberg & Jaeger’s (2016) proposal that the degree and level of predictive pre-activation depend on its expected utility, which is shaped by comprehenders’ goals and their assessment of the reliability of prior knowledge and bottom-up input.

To conclude, comprehenders seem to consider the phonological variability between speakers in predicting the upcoming speech. These findings shed light into both the level(s) of representation and the processes involved in generating predictions. Our results support the notion that linguistic prediction involves the pre-activation of phonological representations. Furthermore, they also suggest that phonological predictions do not derive solely from the passive spreading of activation between higher-level representations that percolate to phonological representations. Rather, it may be influenced by other mechanisms that are more sensitive to the phonological characteristics of the speaker such as event simulation or internal generative models. The predictive nature of the task represents a potential limitation of the present study, as it may have influenced participants’ reliance on phonological predictions. It would be crucial to investigate whether and how phonological predictions are implemented in natural conversations, where additional communicative goals and constraints may shape predictive processes. This would provide a more comprehensive understanding of the flexibility of the perceptual system to predict the upcoming speech.

## Data and code availability

Materials, data and analysis scripts will be made publicly available on OSF upon acceptance of the manuscript.

## Ethics approval

The research adhered to the principles outlined in the Declaration of Helsinki. The research protocol was approved by the Ethics Committee for Psychological Research of the University of Padova (protocol number: 5181). Participants provided their informed consent before participating in the experiment. They also consent to the use of their data (collected anonymously) for research aims.

## CRediT author statement

MS: Conceptualization, Investigation, Data curation, Formal analysis, Visualization, Writing - Original Draft. FV: Conceptualization, Writing - Review & Editing. SG: Conceptualization, Writing - Review & Editing. LC: Investigation, Writing - Review & Editing. FP: Conceptualization, Supervision, Funding acquisition, Writing - Review & Editing.

## Footnotes

1 In the ERP literature both PCA (Principal Component Analysis, see Dien & Frishkoff, 2005; Dien, 2012) and EFA (Exploratory Factor Analysis, see Scharf et al., 2022) have been used to estimate the underlying components of the ERP data. However, the differences between PCA and EFA estimates are negligible for ERP data due to the high correlations between sampling points and to the high number of variables (Dien & Frishkoff, 2005; Scharf & Nestler, 2019; Widaman, 1990, 2007).

